# Deep learning guided discovery of antibacterial polymeric nanoparticles

**DOI:** 10.1101/2025.05.13.653905

**Authors:** Yuhui Wu, Cong Wang, Xintian Shen, Yan Chen, Haiping Wang, Bocheng Xu, Yifeng Chen, Wenbin Dai, Yue Huang, Lingyun Zou, Jian Ji, Peng Zhang

## Abstract

Drug-resistant bacterial infections pose a serious threat to global health, driving the development of novel antibacterial strategies beyond classic antibiotics. Self-assembled polymeric nanoparticles (SANPs) have emerged as promising candidates due to their unique structural properties, which enhance membrane interaction, drug loading, and therapeutic accumulation. The structural diversity and tunable properties of SANPs provide a flexible design platform for antibacterial applications. However, navigating the vast chemical space of SANPs remains a significant challenge due to the complex structure-activity relationships. To address this, we developed PolyCLOVER, a deep learning-guided framework for the efficient discovery of antibacterial SANPs. The framework integrates multi-stage self-supervised learning, active learning, and high-throughput experimentation to iteratively guide the discovery of SANPs with potent antibacterial activity and low toxicity. Through this framework, we identified three potent SANPs within a combinatorial library of ∼100,000 poly(β-amino ester)s, with minimum inhibitory concentrations of 4 μg/mL and 8 μg/mL against clinically multidrug-resistant *S. aureus* and *A. baumannii*, respectively. Moreover, they can also act as antibiotic carriers, restoring the sensitivity of pathogens to penicillin G. *In vivo* studies demonstrated their therapeutic efficacy both as monotherapies and in combination therapies with antibiotics. These results demonstrate PolyCLOVER as a powerful framework for discovering multi-functional SANPs to combat antibiotic resistance.

## Main

Bacterial infections, particularly those caused by drug-resistant strains, pose a significant threat to human health^1,2^. Classic antibiotics penetrate bacterial membranes to target intracellular proteins^3^, whereas bacteria have evolved resistance mechanisms such as efflux pumps^4^, enzymatic degradation^5^, and modification^6^ to evade these treatments^7^. Efforts to tackle bacterial resistance through alternative pathways and mechanisms are gaining attention, with nanoparticles (NPs) emerging as a potent option for new antibacterials or antibiotic adjuvants^3^. NPs offer precise control over charge and hydrophobicity distribution, which enhances their interaction with bacterial cell membranes. This interaction disrupts normal bacterial physiological functions, reducing the likelihood of drug resistance^8-10^. Additionally, NPs facilitate chemical modification, which allows for easy integration with antibiotics and enhances their accumulation within bacterial membranes to counteract drug resistance more effectively^11^. Furthermore, the drug delivery system based on NPs can efficiently transport antibiotics across dense biofilms, thereby overcoming this critical barrier^12^. The unique size and finely engineered surface properties of NPs also enable target delivery of antibacterials to ensure suitable distribution within infected tissues, improving therapeutic outcomes^13,14^.

Designing nanodrug delivery systems that also function as antibacterials requires considerable effort to regulate charge distribution and hydrophilic-hydrophobic balance^15-18^. Polymers stand as a promising platform for assembling these building blocks into NPs through self-assembly (SANPs) driven by noncovalent interactions^19^. However, a significant challenge here lies in developing polymer libraries that support the creation of copolymeric SANPs with a wide range of functional moieties. Poly(β-amino ester)s can be synthesized through the Michael addition reaction between diacrylate and amine monomers, enabling extensive customization of the polymer chemical properties^20^. This step-growth polymerization produces linear polymers with ester and tertiary amine structures in their skeleton^21,22^. Such polymers have been widely explored for their ability to deliver nucleic acids and proteins due to their inherent positive charge^23-25^. Therefore, we envision that the structural diversity of poly(β-amino ester)s will unlock their potential for establishing a versatile library of antibacterial agents.

Handling the vast possibilities within the library of SANPs presents a significant challenge for traditional experimental approaches. Machine learning (ML) has been successfully applied to explore the chemical space of polymers due to its ability to model complex chemical interactions^26-39^. These studies typically train supervised learning models using publicly available labeled data, followed by predicting the properties of a defined pool of candidate polymers. However, existing approaches when applied to antibacterial SANPs design are limited by several key issues: 1) There is a lack of sufficient data to support robust modeling. Despite some prior studies on antibacterial SANPs, these data have limited applicability due to significant differences in material chemistry and antimicrobial characterization. 2) The inherent structural complexity of SANPs hinders effective feature extraction and reduces the expressiveness of material representations, thereby constraining generalization across the vast and diverse chemical space.

Herein, we proposed a comprehensive framework (PolyCLOVER) to solve the above challenges and promote the discovery of multi-functional poly(β-amino ester) SANPs, which serve as antibacterials to combat drug resistance and are also endowed with antibiotic delivery capacity. We designed a poly(β-amino ester) library comprising about 100,000 candidates based on a combinatorial strategy, covering a broad chemical space (**Fig. 1a**). PolyCLOVER nominates SANPs with both high antibacterial activity and low toxicity using multi-stage self-supervised learning, active learning, and high-throughput experiments, achieving efficient navigation in the chemical space. The ensemble multi-task predictor recommends candidate combinations based on properties, diversity, and uncertainty, which are then evaluated through experiments. These data are then used to update the pre-trained model and predict the next round of candidates (**Fig. 1b**). Through the proposed framework, three polymers with outstanding antibacterial efficacy and low hemolytic toxicity were successfully identified. We further validated their self-assembly properties and tested their therapeutic efficacy in a mouse pneumonia model. Meanwhile, the candidate polymer showed high encapsulation efficacy and loading capacity when co-assembled with penicillin G. The co-assembled NPs showed a synergistic antibacterial effect and restored the sensitivity of drug-resistant Gram-positive pathogens to penicillin G, demonstrating a combined therapeutic effect in a mouse peritonitis model (**Fig. 1c**). PolyCLOVER is anticipated to significantly streamline the discovery for SANPs integrated with antibacterial and drug repurposing potential.

**Fig. 1.**
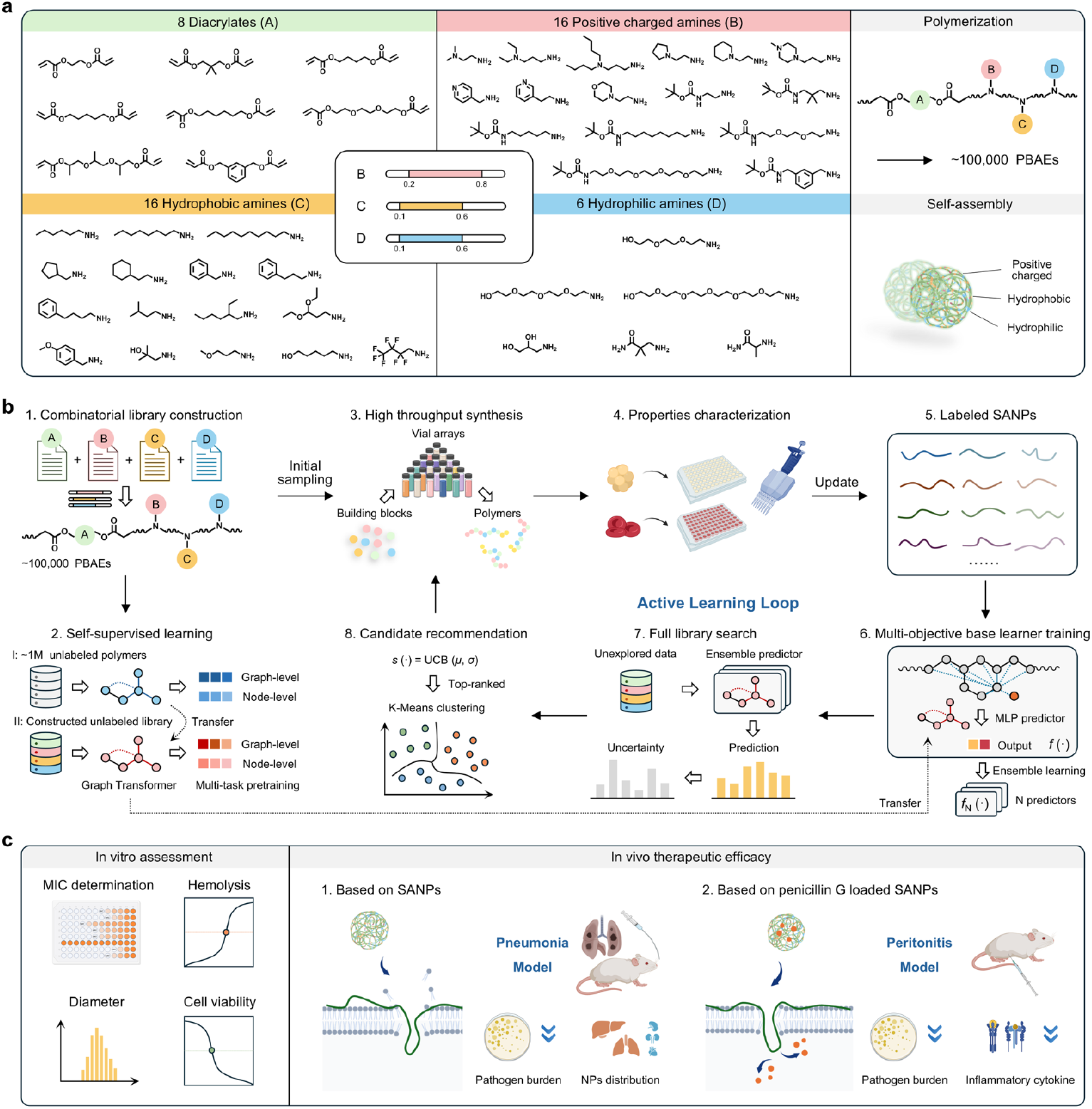
Overview of PolyCLOVER. **a**, Combinatorial library construction. The library was constructed based on Michael addition between diacrylate and amine monomers. 8 diacrylates and 38 amines were selected based on solubility and cost from commercially available sources, including 16 amines with tertiary amine side groups or Boc-protected primary amine groups, 16 with hydrophobic side groups, and 6 with hydrophilic side groups. The ratios of B, C, and D were controlled within appropriate ranges to drive self-assembly, resulting in the formation of ∼100,000 poly(β-amino esters) SANPs. **b**, PolyCLOVER workflow. A graph encoder was pre-trained using a two-stage, multi-task self-supervised learning based on the constructed library. An initial subset of the library was sampled, followed by high-throughput synthesis and characterization of antibacterial activity and hemolytic rate to generate a labeled dataset. The graph encoder was then fine-tuned to form an ensemble predictor consisting of 20 base learners. The predictor estimated the activity and predictive uncertainty of unlabeled SANPs in the library, guiding the iterative selection of new candidates via UCB acquisition and K-Means clustering. This process was iteratively repeated. **c**, In vitro/in vivo validation. Additional in vitro experiments were conducted to characterize the detailed antibacterial activity, cytotoxicity, and particle size of the selected SANPs. The in vivo therapeutic efficacy of the SANPs, either alone or in combination with penicillin G, was validated in pneumonia and peritonitis models.

### Multi-stage self-supervised learning boosts model performance on limited data

Previous studies have demonstrated that positive charges present in polymers can effectively interact with the negatively charged bacterial cell membrane, while groups such as alkyl chains can insert into the bacterial cell membrane and disrupt its normal physiological function, thereby achieving a bactericidal effect^40-42^. Accordingly, positively charged and hydrophobic units were incorporated into polymers to impart potential antibacterial activity. We also introduced hydrophilic units and regulated the ratios of each component in a suitable range to promote the self-assembly of the polymers into stable SANPs. Guided by this design rationale, a combinatorial library comprising 98,304 (∼100,000) poly(β-amino ester) SANPs was constructed via a multi-component Michael addition reaction (**Supplementary Table S1, 2**).

Next, we randomly selected 220 samples from the constructed library for laboratory synthesis and characterization to form the initial dataset. The initial dataset spans a vast chemical space, incorporating all monomer types presented in the library (**Fig. 2a, Supplementary Fig. S1**). We chose Methicillin-resistant *Staphylococcus aureus* (MRSA) as the model pathogen because it stands as one of the most prevalent pathogens, capable of causing life-threatening diseases such as sepsis, pneumonia, and multi-organ failure^43^. The antibacterial activity against MRSA and hemolytic behavior of the samples were quantitatively assessed using optical density (OD) values at 595 nm and the hemolysis ratio, respectively. The results reveal that the probability of identifying effective SANPs through random sampling in the extensive combinatorial library is very low. Most SANPs exhibit weak antibacterial activity, while a few samples with high antibacterial activity show high hemolytic toxicity (**Fig. 2b**). These findings underscore the considerable challenge of achieving a balance between high antibacterial activity and low hemolytic toxicity, highlighting the importance of ML in guiding the design of antibacterial SANPs.

**Fig. 2.**
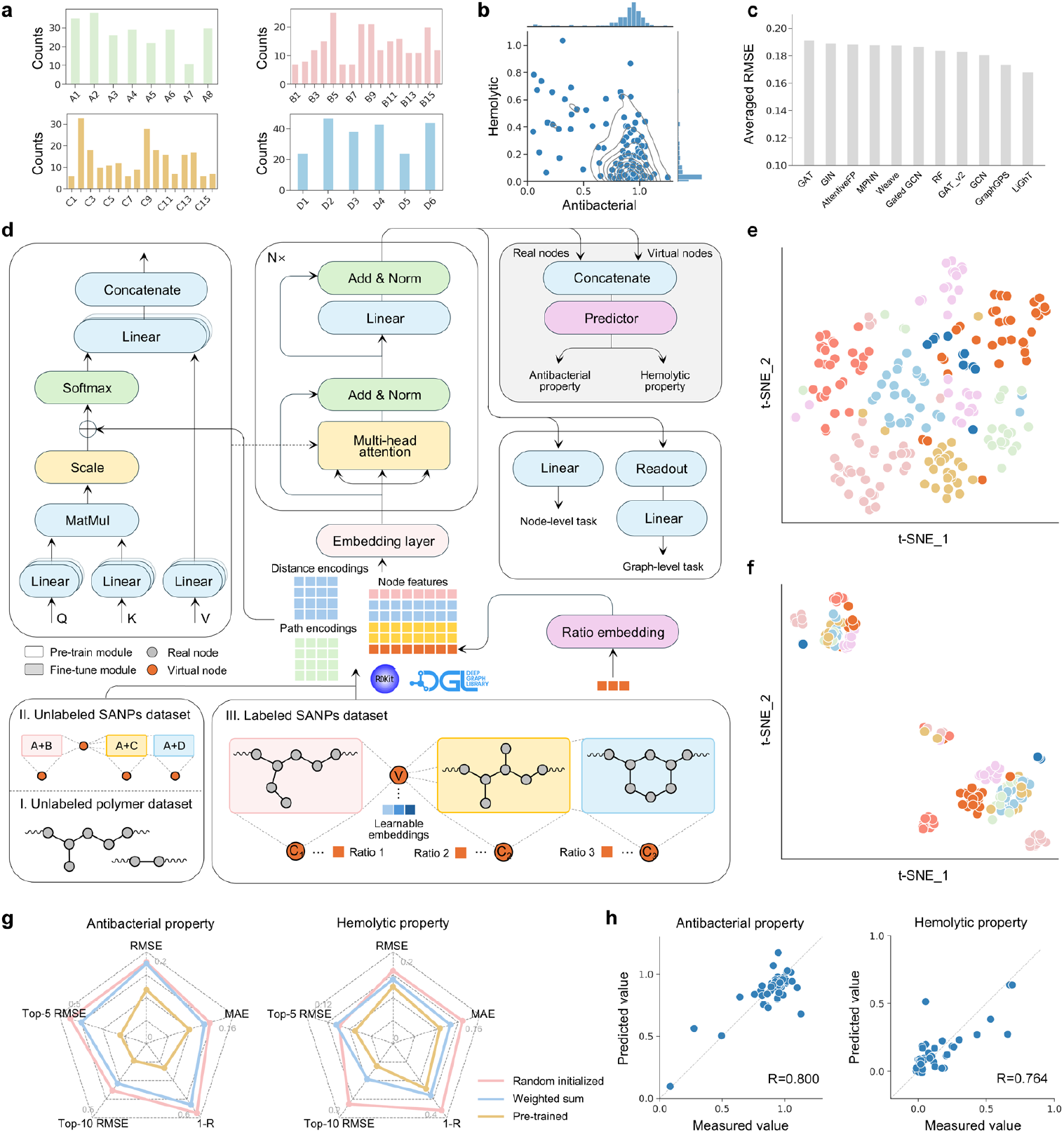
Model training and evaluation. **a**, The number of different types of building blocks in the initial labeled dataset. **b**, The distribution of antibacterial activity and hemolytic property in the initial dataset. Marginal histograms along the top and right axes show the distributions of each property. Contour lines were plotted using kernel density estimation with a threshold of 0.15. **c**, Average test performance of different backbones on antibacterial and hemolytic properties. The mean value of the test root mean square error on three independent runs is reported. **d**, Data preprocessing and model architecture. The Graph Transformer was pre-trained sequentially on an unlabeled polymer dataset and the constructed unlabeled NP dataset to learn rich structural features. It was then fine-tuned using the labeled NP dataset to predict antibacterial and hemolytic properties. **e, f**, t-SNE visualization of representations generated by the pre-trained (**e**) and un-pretrained (**f**) graph encoder. Each data point is colored according to the type of diacrylate used. **g**, Test performance of the proposed approach. Pearson correlation coefficients are shown as 1-R to facilitate visualization on the radar plot, where a smaller enclosed area indicates better performance. The mean value of the test root mean square error on three independent runs is reported. **h**, The measured value versus predicted plots for the proposed model. The dashed line represents the ideal case, where y = x. The Pearson correlation coefficient is shown in the lower right corner.

After collecting the initial dataset, we employed graph neural networks (GNNs) for modeling, due to their strong capability in capturing molecular structural features^44,45^. However, the intrinsic complexity of copolymers, particularly the absence of a well-defined and consistent topology, makes it challenging to represent them using the vanilla molecular graph. To enable effective message passing between subgraphs corresponding to individual building blocks, virtual nodes were introduced as communication bridges, following our previously reported approach^46^. In addition, three local virtual nodes were connected to their respective subgraphs to encode the copolymerization ratios. More details about graph construction are provided in the Methods section. We evaluated the average test performance of 10 commonly used GNN backbones on antibacterial and hemolytic tasks and finally selected a Graph Transformer^47^ with multi-head attention (**Fig. 2c, Supplementary Table S3**).

Self-supervised learning has demonstrated remarkable success across various domains of artificial intelligence by leveraging intrinsic structural information in large-scale unlabeled data, enabling models to learn general representations that can be effectively transferred to downstream tasks^48-51^. This paradigm significantly reduces reliance on manual annotations and improves model generalization. To this end, we employed a multi-task self-supervised learning strategy for the polymer encoder, using a large corpus of unlabeled polymer data. The model is jointly trained on node-level and graph-level objectives to capture hierarchical structural representations of polymers^46^. Notably, our pool-based screening strategy enables the property prediction model to access candidate SANPs that will be evaluated during the inference phase. That is, the model is exposed to the candidate set used during prediction, effectively establishing a transductive learning scenario. For this reason, we propose a two-stage pre-training approach, in which the encoder is further pre-trained on the entire accessible unlabeled candidate pool to fully exploit the structural priors inherent in the screening space and to bridge the gap between pre-training and fine-tuning domains. The ratio information of individual building blocks was extracted using a ratio embedding layer and assigned as the initial features of the corresponding virtual nodes (**Fig. 2d**).

Subsequently, we analyzed the model’s latent space to evaluate the effect of the two-stage pre-training. The representations generated by the pre-trained model and the unpretrained model were visualized using t-SNE, respectively (**Fig. 2e, f, Supplementary Fig. S2-6**). It can be observed that the representations generated by the pre-trained model are evenly distributed in the latent space and exhibit clear clustering patterns, where structurally similar SANPs (the same diacrylates) show high consistency in their representations. In the unpretrained model, the representations are compressed into narrow regions, indicating limited structural awareness and reduced information retention. These results demonstrate that the pre-trained model effectively captures key structural features while preserving maximal information^52^.

The pre-trained model was then fine-tuned on the initial dataset to predict the antibacterial and hemolytic properties of SANPs. We compared the test performance of the two-stage self-supervised learning approach with baseline methods (**Fig. 2g, h, Supplementary Fig. S7, 8**). Due to the extreme scarcity of data, the randomly initialized model fails to capture structural patterns, resulting in poor performance. The weighted sum approach used ratio-weighted building block representations extracted by the pre-trained encoder, followed by a prediction head. Its performance showed a slight improvement but remained inadequate. In contrast, our proposed approach achieves significantly better performance across all metrics, demonstrating its effectiveness under data-limited conditions. Moreover, the fine-tuned latent space reveals an ordered, continuous distribution that aligns with activity levels, reflecting better capture of the underlying structure-activity relationship (**Supplementary Fig. S9, 10**). Therefore, we adopt this model as the base learner for subsequent structure optimization of SANPs.

### PolyCLOVER drives iterative improvements in antibacterial activity and hemocompatibility

Although the model was optimized, the limited amount of training data still poses a significant challenge for generalizing across the full combinatorial library. To address this, PolyCLOVER constructs an ensemble predictor composed of 20 homogeneous base learners, which is then embedded into a Bayesian optimization-based active learning framework. This pipeline facilitates efficient exploration of the vast chemical design space of SANPs to identify structures with potentially optimal properties.

Then, the ensemble predictor was employed to predict the activities of unlabeled SANPs in the remaining combinatorial library. The mean prediction across base learners was used as the activity estimate, while the standard deviation served as a measure of predictive uncertainty. Each candidate was then scored using the Upper Confidence Bound (UCB) acquisition function, which balances exploration of uncertain regions with exploitation of potentially optimal candidates. To further improve optimization efficiency, we applied the K-means algorithm to cluster the top-ranked SANPs and selected representative structures from each cluster to enhance data diversity. PolyCLOVER then recommended 60 candidate SANPs for wet-lab synthesis and characterization. The newly generated data were subsequently incorporated into the initial dataset to support the next round of optimization **(Fig. 3a**). More details can be found in the Methods section.

**Fig. 3.**
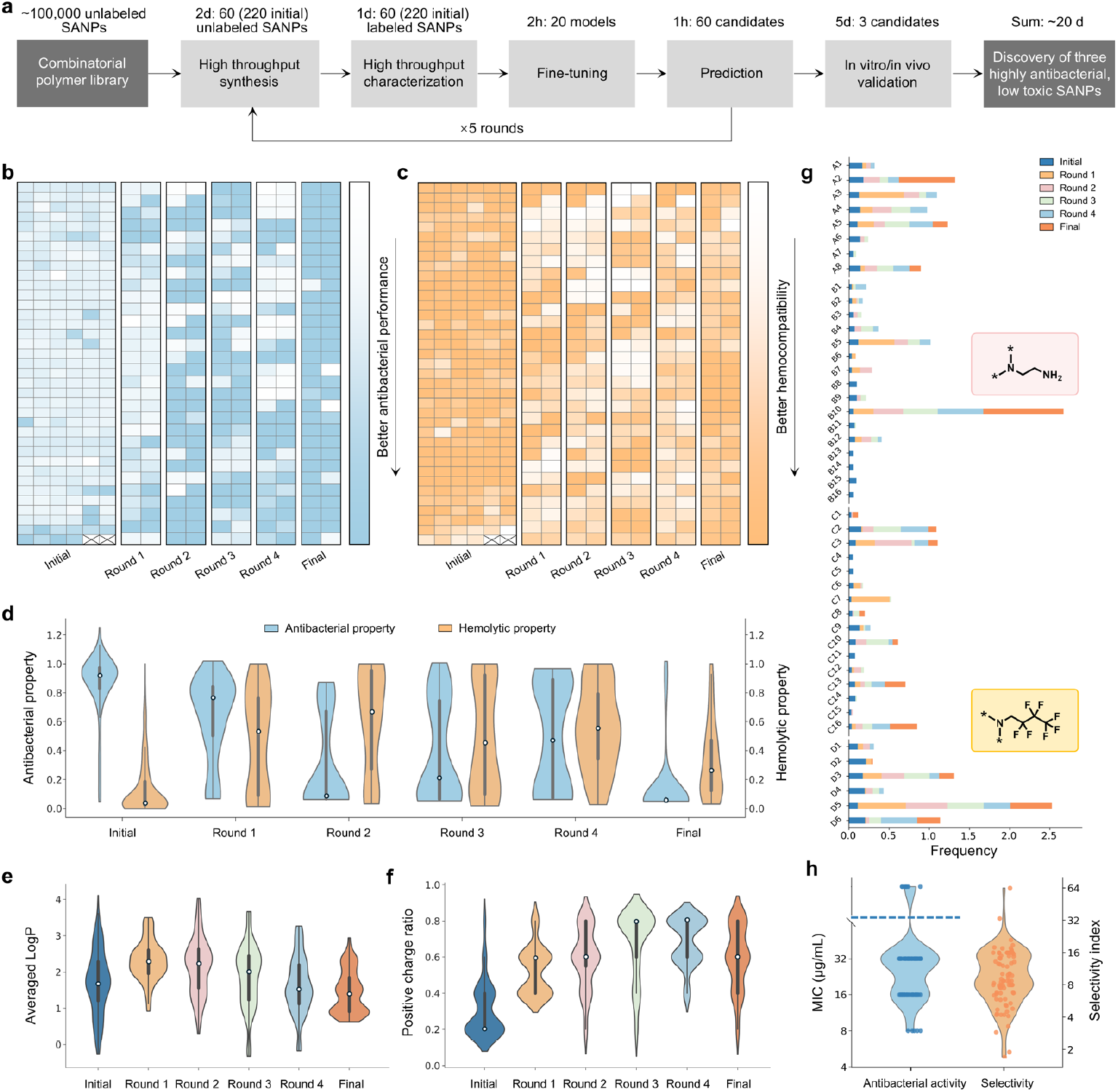
Bayesian optimization-based active learning process. **a**, Time spent in each step of PolyCLOVER for identified SANPs screening. **b, c**, Heat maps of antibacterial (**b**) and hemolytic (**c**) properties in each round. **d**, Density plots of antibacterial and hemolytic properties in each round. Lower values represent better antibacterial activity and improved hemocompatibility. The white dot in each data group represents the median value. **e, f**, The evolution of average LogP (**e**) and positive charge ratio (**f**) over iterative optimization rounds. The LogP value of each NP is calculated as a weighted sum of the LogP values of its building blocks according to their compositional ratios. **g**, The frequency of individual building blocks over iterative optimization rounds. Boxed insets show the structures of B10 and C16. **h**, The MIC value and selectivity index of the SANPs that meet the criterion. Samples above the dashed line represent MIC greater than 32 µg/mL. The selectivity index is calculated as the ratio of HC_50_ to MIC.

After four rounds of iterative optimization, the activities of the recommended SANPs gradually stabilized, suggesting convergence of the learning process (**Fig. 3b, c, d, Supplementary Fig. S11-15**). At this stage, a greedy acquisition strategy was employed to further exploit the most promising candidates. Compared to the initial dataset, the antibacterial activity of the selected SANPs showed a substantial improvement. Although the hemolytic rate increased slightly in the intermediate rounds due to the enhancement of antibacterial potency, it was successfully reduced to an acceptable level in the final round of optimization. As a result, a substantial number of SANPs identified in the final round exhibited both high antibacterial efficacy and low hemolytic toxicity.

To gain deeper insight into the model’s behavior, we further investigated the evolution of chemical structures and key physicochemical properties of the SANPs recommended by PolyCLOVER. In the first round, an increase in LogP indicated a shift toward higher hydrophobicity, suggesting that the model initially prioritized antibacterial activity. However, excessive hydrophobicity (e.g., from longer alkyl chains) was often associated with a higher potential for cell toxicity^10^, the dual-objective task subsequently regulated this property to balance antibacterial activity and biocompatibility (**Fig. 3e**). Additionally, the initially low ratio of positive charges, which may have limited interactions with negatively charged bacterial membranes, was substantially increased during optimization (**Fig. 3f**). A sharp increase in the de-protected primary amine of B10 suggested that the model favored a short chain length of positive charge (**Fig. 3g**). We also found the unique integration of polyfluorocarbon unit (C16) effectively preserved antibacterial efficacy while reduced toxicity, which aligned with previous studies^53^.

To further identify SANPs, we established a preliminary criterion for high-potential candidates: an antibacterial optical density (OD) value of less than 0.15 and a hemolytic rate of less than 50%. All SANPs meeting this criterion were synthesized and purified in a single batch, followed by detailed evaluation of their antibacterial and hemolytic properties. The minimum inhibitory concentration (MIC) and half-hemolytic concentration (HC_50_) were determined for each polymer, and the selectivity index was calculated as the ratio of HC_50_ to MIC (**Fig. 3h, Supplementary Table S4**). The results showed that most SANPs exhibited strong antibacterial activity and low hemolytic toxicity, with selectivity indices exceeding eight. Finally, samples with a selectivity index greater than 20 were selected, resulting in the identification of three optimal SANPs. Notably, PolyCLOVER enabled the discovery of three highly antibacterial and low-toxic SANPs from a combinatorial library of ∼100,000 polymer candidates in just 20 days, demonstrating its remarkable efficiency in accelerating the discovery of new functional materials within large design spaces.

### Identified SANPs exhibit promising *in vitro* performance

The identified SANPs were designated as H1, H2, and H3 **(Fig. 4a)**. The positively charged segments of the SANPs consisted of deprotected primary amines with short chains. The hydrophobic portions featured either a relatively short tertiary alcohol or a short chain with fluorinated side groups, which likely mitigated the hemolytic toxicity associated with longer hydrophobic chains. The hydrophilic components included oligomeric polyethylene glycol (PEG) side groups in H1 and H3, while H2 contained a unique amide side group. Notably, the proportion of hydrophilic components was the lowest among the three segments, while the proportion of positively charged groups was the highest, which is crucial for interaction with negatively charged bacterial cell membranes. Additionally, due to an excess of diacrylate during polymer preparation, all three SANPs retained double bond residues at both ends, allowing for potential future functional modifications (**Supplementary Fig. S16**). The MIC of all three SANPs were 8 μg/mL. Both the HC_50_ (H1 = 168 μg/mL, H2 > 512 μg/mL, H3 = 266 μg/mL) and the half-maximal inhibitory concentration (IC_50_; H1 = 53 μg/mL, H2 = 162 μg/mL, H3 = 109 μg/mL) followed a consistent trend, with H2 exhibiting the lowest toxicity among the three. (**Supplementary Fig. S17**). These findings underscore the necessity of leveraging a dual-objective optimization strategy to achieve a balance between antibacterial efficacy and toxicity. The self-assembly properties of H1-3 were confirmed by dynamic light scattering (DLS) and transmission electron microscopy (TEM). Three SANPs exhibited particle diameters of approximately 100 nm, with H2 having a larger diameter compared to H1 and H3, and all three SANPs carried positive charges **(Fig. 4b)**.

**Fig. 4.**
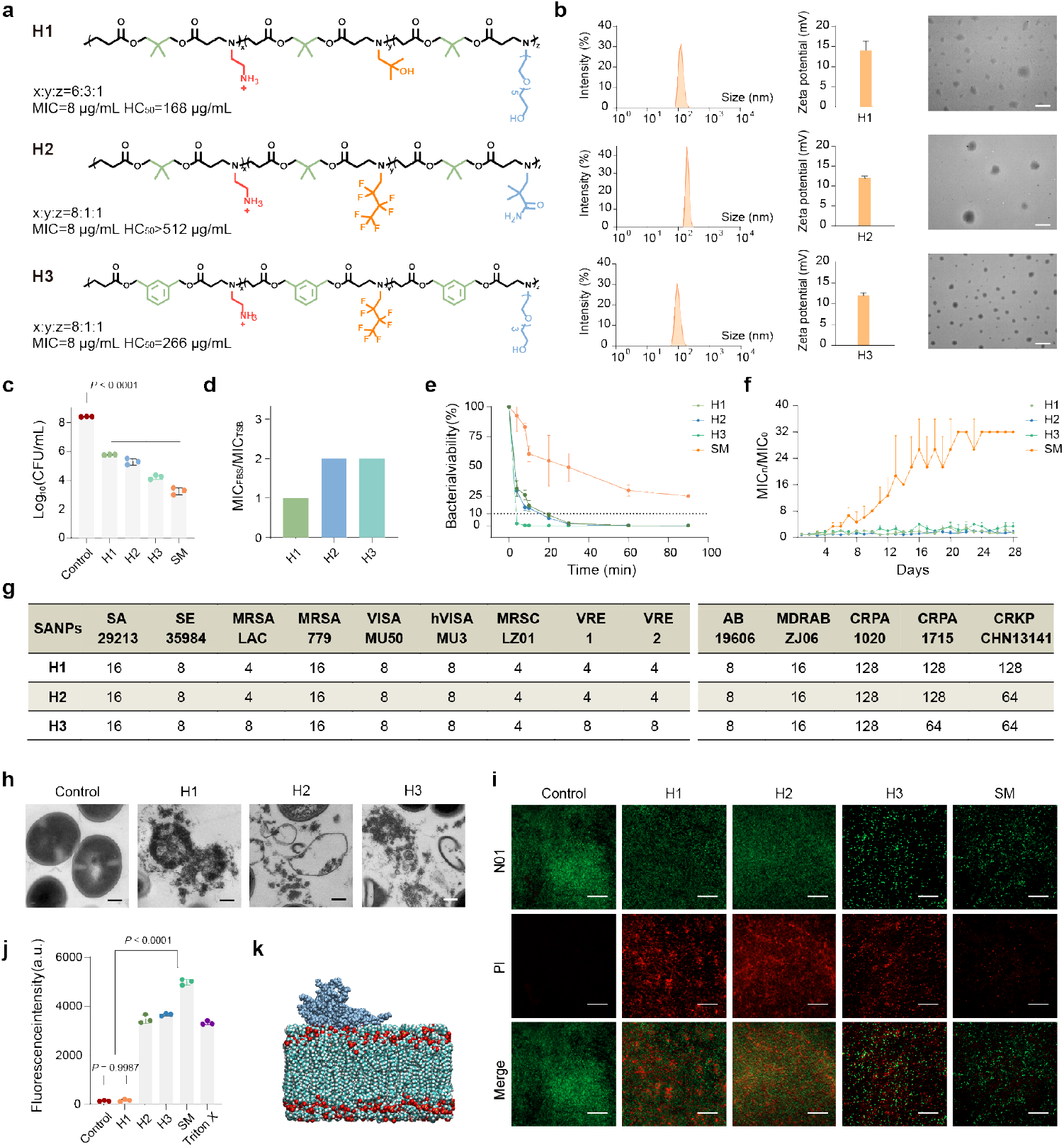
Structures, in vitro antibacterial activity, and antibacterial mechanism of identified SANPs. **a**, Detailed information of the selected identified SANPs (H1, H2, H3), including molecular structure, monomer ratio, MIC values, and hemolytic toxicity. **b**, Size distribution, zeta potentials, and TEM images of identified SANPs. Scale bar: 1 μm. **c**, Bactericidal performance of SANPs and SM at 32 µg/mL for 9 h incubation. Statistical analysis was performed using one-way ANOVA. **d**, Changes in MIC values of identified SANPs in Tryptic Soy Broth (TSB) and Fetal Bovine Serum (FBS). **e**, Bactericidal kinetics of identified SANPs and SM at 32 µg/mL within 100 minutes. **f**, Evolution of resistance to identified SANPs and SM in MRSA after 28 days of passaging in TSB media. **g**, MIC values (µg/mL) of the identified SANPs against 3 reference strains (*S. aureus* ATCC29213, *S. epidermidis* ATCC35984, and A. baumannii ATCC19606) and 11 clinically isolated drug-resistant pathogens (*S. aureus*-MRSA, VISA, *S. capitis*-MRSC, *E. faecalis*-VRE, *A. baumannii*-MDRAB, *P. aeruginosa*-CRPA, *K. pneumoniae*-CRKP). Gram-positive and Gram-negative bacteria are shown on the left and right, respectively. Each test was performed in three independent experiments. **h**, TEM images of MRSA after incubation with identified SANPs for 10 min at 64 µg/mL. Scale bar: 200 nm. **i**, Live/dead bacterial staining of MRSA after incubation with identified SANPs for 15 min at 64 µg/mL. Scale bar: 20 μm. **j**, Fluorescence analysis of membrane potential disruption induced by identified SANPs and SM at 64 µg/mL. Statistical analysis was conducted using one-way ANOVA. **k**, Molecular dynamics simulation of the interaction between H2 and the cell membrane. In panels **b, h**, and **i**, experiments were performed in triplicate, yielding consistent results, and a representative image is shown. In panels **b, c, e, f**, and **j**, data are shown as mean ± s.d. (n = 3 independent replicates).

We further evaluated the bactericidal efficacy of the identified SANPs by benchmarking them against streptomycin (SM), a broad-spectrum antibiotic. After 9 hours of co-incubation at 32 μg/mL, H1 and H2 reduced MRSA viability by 3 orders of magnitude, while H3 achieved a 4-order reduction (**Fig. 4c**). SM exhibited the strongest antibacterial effect, with a 5-order reduction. Meanwhile, when the tryptic soy broth (TSB) medium was replaced with fetal bovine serum (FBS), a more complex environment, we found that the SANPs maintained their antibacterial efficacy, with MIC values fluctuating by only 1-2 fold. This stability may be attributed to the biologically inert fluorinated and PEG side chains **(Fig. 4d)**. Next, we assessed the bactericidal kinetics of the SANPs, which is another key metric in the practical application of antibacterials. (**Fig. 4e)**. All three SANPs demonstrated rapid bactericidal action. Notably, H1 and H2 eliminated 90% of bacteria within 60 minutes, while H3 achieved this reduction in just 4 minutes. In contrast, SM exhibited significantly slower killing kinetics. As an antibiotic that inhibits bacterial protein synthesis by targeting ribosomes^54^, SM requires prolonged exposure, aligning with prior reports that antibiotics generally exert bactericidal effects over several hours or longer^55^. These findings highlighted that the identified SANPs not only outperformed antibiotics in bactericidal speed but also likely operated via a different antibacterial mechanism. Furthermore, compared to SM, the SANPs exhibited a substantially lower tendency for resistance development. Over a 28-day testing period, the MIC values for H1-3 fluctuated modestly (1-4 fold), while SM showed a significant 28-fold increase in MIC (**Fig. 4f**). Among 11 clinically isolated drug-resistant pathogens, the SANPs exhibited antibacterial efficacy comparable to that observed in reference strains. In particular, the SANPs exhibited potent activity against all tested Gram-positive bacteria, whereas activity among Gram-negative strains was restricted to A. baumannii. These results indicate that the antibacterial spectrum of the SANPs is selective and biased towards Gram-positive pathogens. (**Fig. 4g**).

Subsequently, we investigated the antibacterial mechanism of the identified SANPs. Transmission electron microscopy (TEM) images were obtained after 5 minutes of co-incubation with the identified SANPs (**Fig. 4h**). In the control group, the MRSA membrane maintained a clear and intact structure. In contrast, after interaction with the SANPs, significant membrane disintegration was observed, leading to the release of bacterial cell contents and resulting in bacterial death. Live/dead bacterial staining assays were performed after 5 minutes of exposure to SANPs H1-3 (**Fig. 4i**). The assays employed N01, which penetrated both live and dead bacteria, binding to nucleic acids and emitting green fluorescence, while propidium iodide (PI) entered only damaged or dead bacteria, binding to DNA and emitting red fluorescence. Confocal fluorescence microscopy revealed intense red fluorescence in MRSA exposed to SANPs H1-3, even after a short incubation period. To assess their effect on bacterial membrane potential, we used the fluorescent probe DiSC_3_(5) (**Fig. 4j**). This probe accumulates in intact bacterial membranes, leading to fluorescence quenching. When the membrane is disrupted, the probe leaks from the membrane, causing fluorescence emission as its concentration decreases. Bacterial suspensions without SANPs exhibited negligible fluorescence intensity. In contrast, the addition of Triton X-100, a strong surfactant, resulted in a significant increase in fluorescence emission due to membrane destabilization. SANPs H1-3 induced robust fluorescence signals, confirming their ability to disrupt bacterial membrane potential. SM, however, did not induce fluorescence, consistent with its non-membrane-targeting antibacterial mechanism. The membrane-attaching mechanism of the SANPs was further revealed by molecular simulations. All-atom molecular dynamics results demonstrate that when the SANPs approach the cell membrane, they rapidly adsorb to the surface. This process is primarily driven by electrostatic interactions. (**Fig. 4k, Supplementary Fig. S18**). Analysis of these results suggests that the identified SANPs likely exert their bactericidal effects through strong interactions with the cell membranes of MRSA in a short timeframe.

### Identified SANPs eradicate lung infection with high efficacy and biocompatibility

The therapeutic efficacy of the identified SANPs was further investigated in a mouse lung infection model. MRSA-induced acute pneumonia is among the most encountered lung infections, featuring a high incidence rate and significant mortality^56^. Female mice were inoculated with MRSA to establish an acute pneumonia model by intratracheal instillation (**Fig. 5a**). The identified SANPs were administered via tail intravenous injection (IV injection) on the following day. Mice were euthanized one day post-treatment, and lung tissues were collected for morphological observation and bacterial counting. The PBS-treated group presented unhealthy dark red lung appearances. In contrast, the lungs of the identified SANPs-treated group appeared healthy and pink. Bacterial counts from tissue homogenates indicated that the SANPs-treated group demonstrated significant improvement, reducing bacterial counts by three orders of magnitude, comparable to the results observed in the SM-treated group (**Fig. 5b, c**). The results of H&E staining revealed that the alveolar structure was collapsed in the PBS treatment group. In contrast, it remained porous in the SANPs-treated group, indicating the relief of lung infection. Additionally, Masson staining showed no significant blue fiber deposition, further suggesting that H1-3 exhibited a favorable therapeutic effect (**Fig. 5d**).

**Fig. 5.**
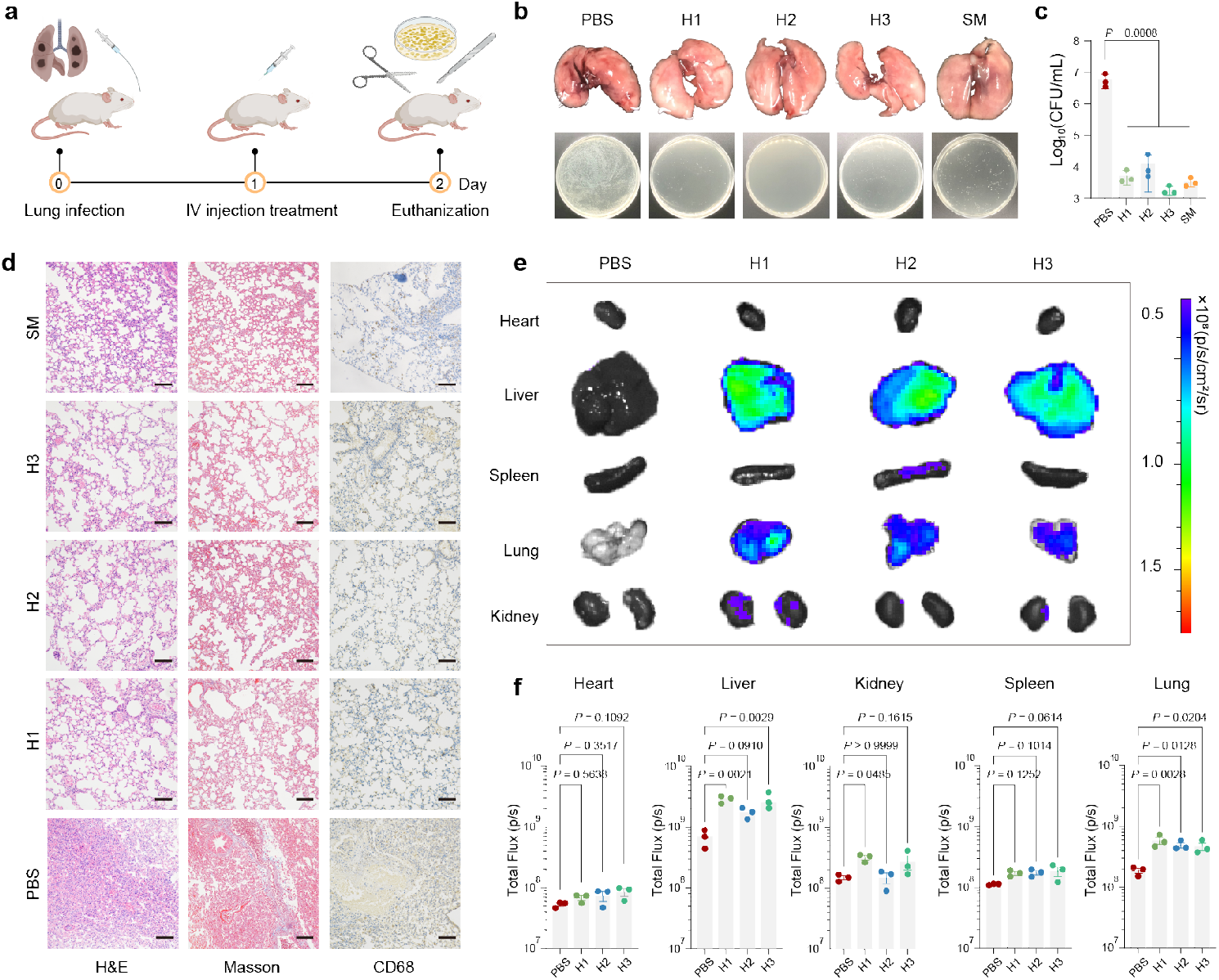
Therapeutic efficacy of identified SANPs in treating acute pneumonia in vivo. **a**, Schematic illustration of the acute pneumonia model construction and treatment protocol in mice. On day 0, mice were anesthetized with isoflurane, and MRSA suspension (10^8^ CFU/mL, 60 μL) was administered intratracheally to induce infection. On day 1, the identified polymers H1, H2, H3 and SM were administered via tail vein injection at a dose of 5 mg/kg. On day 2, the mice were euthanized, and their lungs were excised for morphological observation. The lung tissue was homogenized for further bacterial colony counts. **b**, Visual examination of mouse lung appearance and bacterial colony culture from tissue homogenate to assess infection. **c**, Quantification of lung bacterial load through colony counting in infected mice. **d**, Histological analysis of lung tissues using H&E and Masson staining to evaluate the extent of tissue damage and fibrosis. Scale bar: 300 nm. **e, f**, Quantitative results and fluorescence imaging of the distribution of SANPs within the mice to determine tissue targeting efficiency. In panels **b** and **d**, experiments were performed in triplicate, with consistent results, and one representative image is shown. In panels **c** and **f**, data are expressed as mean ± s.d. (n = 3 independent replicates). Statistical analysis was performed using one-way ANOVA.

To examine the distribution of the SANPs in mice post-injection, rhodamine B was conjugated to the SANPs through a reaction between isothiocyanate and the amine groups (**Supplementary Fig. S19**). Statistical results and fluorescence images of mouse organs were captured 2 hours post-injection (**Fig. 5e, f**). The results revealed that most of the fluorescence was concentrated in the liver, with a notable presence in the lungs, indicating that the SANPs effectively reached lung tissue via the tail vein injection. Next, we assessed the tissue toxicity of the SANPs (**Supplementary Fig. S20**). Histological analysis of heart, liver, spleen, lung, and kidney samples from SANPs-treated mice revealed no significant tissue necrosis compared to the PBS control group. The lung sections exhibited a healthy, loose alveolar structure, and no damage was observed in the glomeruli or renal tubules of the kidney sections. In conclusion, the identified SANPs demonstrated no evident tissue toxicity.

### Identified SANPs restore MRSA susceptibility to penicillin G

The above investigation demonstrated the intrinsic antibacterial activity and self-assembly properties of the identified SANPs. Furthermore, we evaluated the potential of the SANPs as antibiotic adjuvants. It was anticipated that abundant functional groups within the SANPs could be integrated with antibiotics to form nanocomposites, thus enhancing the penetration of antibiotics through bacterial membranes. MRSA resistance primarily arises from the acquisition of the mecA gene, which encodes penicillin-binding protein 2a (PBP2a)^57^. PBP2a has a significantly reduced affinity for β-lactam antibiotics, such as penicillin G (PG), making these agents largely ineffective against MRSA. The integrated nanocomposites held promise for bypassing this resistance barrier by facilitating more effective antibiotic delivery.

We demonstrated that the SANPs efficiently encapsulated PG, thereby reinstating the susceptibility of MRSA to this antibiotic. Checkerboard assays revealed a clear synergistic effect between the identified SANPs and PG, with all Fractional Inhibitory Concentration Index (FICI) values less than 1. Notably, the FICI value between H2 and PG reached 0.375. At sub-MIC concentrations of H2, the MIC of PG was reduced from 128 μg/mL to 4 μg/mL (**Fig. 6a**). At optimal ratios, the SANPs and PG formed stable spherical nanocomposites with diameters ranging from approximately 100 to 200 nm. Transmission electron microscopy (TEM) images obtained in the dry state clearly showed PG crystals encapsulated within the nanocomposites (**Fig. 6b**). Interestingly, in contrast to previous observations, the H2-PG nanocomposites exhibited the smallest size and a more uniform distribution among the three (**Supplementary Fig. S21**). Furthermore, H2 almost completely encapsulated PG, with a drug loading rate close to 50%, indicating a strong interaction between the two components (**Supplementary Fig. S22**). Molecular dynamics simulations further confirmed the presence of electrostatic interactions between H2 and PG, with a distinct assembly trend observed over 50 ns (**Fig. 6c, Supplementary Fig. S23**). These results suggest that, due to the cationic and hydrophobic properties of the polymer, the nanocomposites preferentially accumulate and integrate into bacterial membranes. This leads to passive enrichment of PG at the membrane interface, thereby enhancing its affinity for penicillin-binding proteins (PBPs) and overcoming the low-affinity resistance mechanism. For subsequent *in vivo* evaluations, we selected the H2-PG nanocomposite, which exhibited optimal synergy (FICI = 0.375), the highest encapsulation efficiency (EE = 97%), and a smaller particle size (106 nm).

**Fig. 6.**
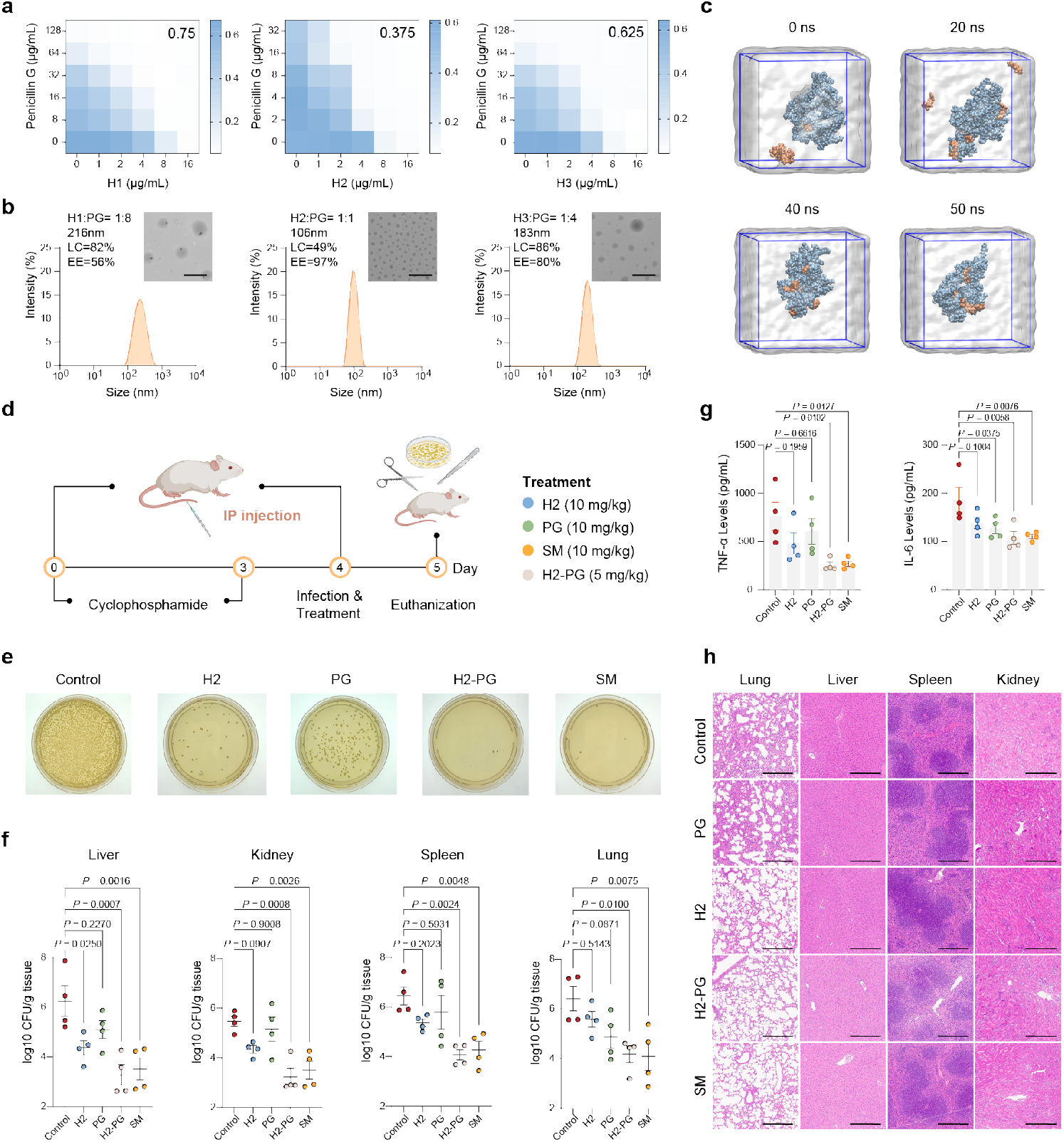
In vitro mechanism validation and in vivo efficacy evaluation of identified SANPs in restoring MRSA susceptibility to penicillin G. **a**, Checkerboard assays assessing the antibacterial synergy between identified SANPs and penicillin G (PG) against MRSA. FICI values are indicated in the upper right corner. **b**, Characterization of SANPs-PG nanocomposites, including size distribution (by DLS), morphology (by TEM; scale bar: 1 μm), encapsulation efficiency (EE%), and loading capacity (LC%) determined by UV-Vis spectroscopy. **c**, Molecular dynamics simulation of the assembly process of H2 and PG. **d**, Schematic illustration of the lethal MRSA peritonitis model and corresponding treatment strategy. Cyclophosphamide was administered intraperitoneally (150 mg/kg on day 1 and 100 mg/kg on day 3) to induce immunosuppression. On day 4, mice were intraperitoneally infected with 1 × 10^7^ CFU of MRSA. Post-infection, mice were treated intraperitoneally with H2 (10 mg/kg), PG (10 mg/kg), SM (10 mg/kg), H2-PG nanocomplex (5 mg/kg), or PBS (100 μL, control). After 24 hours, the mice were euthanized, and samples of serum, spleen, lung, liver, and kidney were collected. **e**, Plate culture images of bacterial colonies obtained from liver tissue homogenates. **f**, Quantification of bacterial loads in liver, kidney, spleen, and lung tissues. **g**, Concentrations of inflammatory cytokines in mouse serum. **h**, Histological analysis of liver, kidney, spleen, and lung sections stained with H&E. Scale bar: 300 nm. In panels **b, e**, and **h**, experiments were performed in triplicates with consistent results, and one representative image is shown. In panels **f** and **g**, data are expressed as mean ± s.d. (n = 4 independent replicates). Statistical analysis was performed using one-way ANOVA.

To evaluate the in vivo therapeutic potential of H2 as a penicillin enhancer, we tested H2-PG nanocomposites in a lethal MRSA peritonitis model, comparing the results with treatments using H2 and PG alone. This model induces systemic diseases, such as sepsis, more readily than respiratory infection models, making it a stringent test for therapeutic strategies. We used a neutropenic mouse model, which is well-established for evaluating antibiotic efficacy^58^. Immunosuppression in mice was induced by cyclophosphamide injection prior to infection. On day 4, mice were infected via intraperitoneal injection with MRSA (1 × 10^7^ CFU), using a lab-induced strain highly resistant to PG *in vitro* (MIC > 64 μg/mL), followed by the administration of various treatments. After 24 hours, the mice were euthanized. Blood and tissues (liver, kidney, spleen, and lung) were harvested for bacterial quantification (**Fig. 6d**). Treatment with H2 (10 mg/kg), PG (10 mg/kg), SM (10 mg/kg), and H2-PG nanocomposites (5 mg/kg) resulted in a reduction of the average bacterial loads in these organs by 1.2, 0.9, 2.3, and 2.5 log units, respectively, compared to untreated controls (**Fig. 6e, f**). This result suggests that in the context of MRSA infection, PG exhibits minimal therapeutic efficacy. In contrast, the H2-PG nanocomposites exhibited anti-infective efficacy comparable to that of streptomycin. Serum inflammatory cytokine analysis revealed a similar trend, with significant reductions in pro-inflammatory markers observed in the H2-PG nanocomposite-treated group (**Fig. 6g**). Histological examination through H&E staining further confirmed that the H2, SM, and H2-PG treatment groups significantly ameliorated tissue damage compared to untreated controls (**Fig. 6h**). These results collectively demonstrate that H2 has substantial potential as an antibiotic adjuvant, capable of restoring bacterial susceptibility to antibiotics.

## Conclusion

In summary, we develop PolyCLOVER, a deep learning-guided framework for the discovery of antibacterial poly(β-amino ester) SANPs. By integrating multi-stage self-supervised learning, active learning, and high-throughput experimentation, PolyCLOVER enables efficient navigation of a vast polymeric chemical space. Within 20 days, the framework successfully identified three lead SANPs from a library of ∼100,000 candidates. These SANPs exhibited strong antibacterial activity against drug-resistant pathogens while maintaining low hemolytic toxicity. Remarkably, these three SANPs not only function as stand-alone antibacterials but also possess antibiotic delivery capacity that restores bacterial susceptibility to antibiotics, as confirmed by both *in vitro* and *in vivo* validation. These results demonstrate the therapeutic potential of the identified SANPs in combating drug-resistant bacterial infections.

More importantly, we established PolyCLOVER as a generalizable and scalable platform applicable to the accelerated design of polymers with diverse functionalities. Through continuous interaction with internal experimental data, it enables autonomous evolution of functional polymers within a large chemical space, achieving complex design objectives without relying on external data. As such, PolyCLOVER offers a blueprint for next-generation polymeric material discovery frameworks that are adaptive, continuously learning, and extensible to the design of materials with precisely tailored functions.

## Methods

### Construction of SANPs library

Poly(β-amino ester)s were synthesized via Michael addition between diacrylates and amines. 8 diacrylates with distinct structures were selected from commercially available sources based on solubility and cost considerations. 16 amines with tertiary amine side groups or Boc-protected amino groups were selected as the positively charged components. Additionally, 16 amines with hydrophobic side groups and 6 with hydrophilic side groups were prepared (Supplementary Table S1). 8 distinct feeding ratios were tested to evaluate the effect of group ratios on the polymer’s antibacterial and hemolytic activity. Eight different feeding ratios were set to ensure its self-assembly capacity (Supplementary Table S2).

### Chemical synthesis of poly(β-amino ester)s

The overall ratio of diacrylate to amine was set at 1.2:1. Initially, the amine monomer solution (3 M in DMF) was pipetted to a reaction vial containing 48 μL of diacrylate (3 M in DMF) according to the specified ratio. The reaction mixture was then heated to 90°C and stirred magnetically for 24 hours. For combinations with Boc-protected amine monomers, 80 μL of trifluoroacetic acid was added after polymerization. The solution was stirred overnight at room temperature to remove the protective group. Finally, the crude product was purified by precipitating in ether/dichloromethane (1:1 in vol%) solution three times.

### Antibacterial and hemolytic activity labelling

The synthesized and purified polymer was dissolved in DMSO at the concentration of 1.28 mg/mL. Fresh methicillin-resistant *Staphylococcus aureus* (MRSA, ATCC 43300) was incubated in Tryptic Soy Broth (TSB) overnight (120 rpm, 37°C). The bacterial suspension was then diluted and calibrated to a concentration of 10^5^ CFU/mL. 195 μL bacterial solution was added to each well in the 96-well plate, followed by addition of 5 μL polymer solution (final concentration at 32 μg/mL). The bacterial suspension was further incubated at 37°C for 9 h. OD values at 595 nm of each well were by the microplate reader.

Hemolysis test was applied to evaluate cytotoxicity. Fresh New Zealand white rabbit blood was collected and centrifuged at 4°C (1500 rpm, 10 min) to remove the supernatant. The remaining red blood cells were resuspended in PBS to obtain 4% red blood cell suspension. 195 μL suspension was added to each well of the 96-well plate, and then the polymer solution of 5 μL was added to achieve a final polymer concentration of 128 μg/mL. The negative control group was added with 5 μL PBS, and the positive control group was added with 5 μL 0.4% Triton X-100. The plates were incubated at 37°C for 1 h. After incubation, the plates were centrifuged at 4°C (1500 rpm, 10 min). After centrifugation, the 100 μL supernatant was gently transferred into a new 96-well plate. OD values at 576 nm was measured, and the hemolysis ratio was calculated according to the following formula:

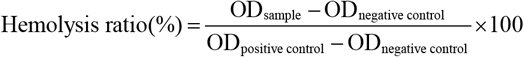

### Model training

In this study, the four-component Michael addition polycondensation product was treated as a random arrangement of three building blocks. The structure of each building block was generated by the reaction module of the RDKit package^59^ according to the general formulas:

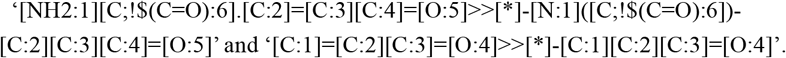

To further promote the model to learn the interactions between components, a global virtual node is introduced here, which is connected to all physical nodes of each component and acts as a bridge for message passing. In addition, the copolymerization ratio of each building block was also introduced in the form of three local virtual nodes, which were connected to all the physical nodes of the corresponding building block, thus forming an ensembled graph^46^. Rdkit was used to calculate the features of atoms and bonds, Mordred^60^ was used to calculate the physicochemical properties of molecules, and DGL package^61,62^ was used to construct the graph *G*.

We selected Graph Transformer (GT) architecture as the backbone of neural network, specifically adopting LiGhT^47^, due to its superior average test performance among ten commonly used GNN variants. Prior to task-specific training, the graph encoder *f*(·) was pretrained in a self-supervised manner using a multi-task learning scheme at both the node and graph levels, which we proposed in our previous work^46^. By jointly optimizing node-level objective and graph-level objective, this strategy allows the encoder to learn multi-scale representations, capturing both fine-grained atomic structure and the broader information of the polymer architecture.

To fully exploit the structural priors inherent in the screening space and to bridge the gap between pre-training and fine-tuning domains, the graph encoder *f*(·) was further pre-trained on the constructed unlabeled library of SANPs. At the node level, we adopted a Masked Node Modeling (MNM) objective ℒ_node_ , where node features were randomly masked, and the model was trained to reconstruct them based on their surrounding context^63^. At the graph level, we employed a contrastive learning objective ℒ_graph_ , where two stochastic masked augmentations *G* and 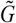 of the same SANPs were treated as a positive pair, while other SANPs in the batch were treated as negatives^64^. The encoder was trained to maximize agreement between representations of the positive pair using InfoNCE loss^65^. The final pre-training loss combines both objectives:

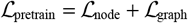

After pre-training, the graph encoder *f*(·) was fine-tuned in a multi-task learning setting. A randomly initialized multilayer perceptron (MLP) projection head was appended to the graph encoder *f*(·) to predict antibacterial and hemolytic activity. The model was trained end-to-end using the Adam optimizer with a batch size of 32. Hyperparameters, including learning rate, dropout, and weight decay, were selected via grid search based on validation performance (**Supplementary Table S7**). MSE loss was applied to each task, and the total loss was computed as their sum. Early stopping with a patience of 20 epochs was employed based on validation performance. All experiments were implemented in PyTorch with DGL and trained on NVIDIA RTX 4090 GPUs.

### Active learning

An active learning framework was applied over multiple rounds to optimize antibacterial and hemolytic properties of SANPs, formulated as a dual-objective regression task. The optimization target for each sample was defined as a scalar score, computed as a weighted sum of the predicted antibacterial activity and hemolysis ratio:

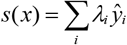

where *λ*_*i*_ is the weighting coefficient that balances the relative importance of the two objectives. *ŷ*_*i*_ is the prediction for the *i*-th task. In this work, *λ* for antibacterial activity was empirically set to 0.7, while *λ* for hemolytic activity was set to 0.3. To estimate the model’s prediction *μ*(*x*) and uncertainty *σ*(*x*), an ensemble learning approach was employed. Specifically, 20 independent models were trained, and for each sample, the predicted mean and standard deviation were calculated from the outputs of the ensemble members. To select candidate SANPs in each round, the Upper Confidence Bound (UCB) acquisition function was employed, balancing exploitation of high-performing samples and exploration of uncertain regions. The acquisition function was defined as:

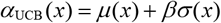

where *β* is a hyper-parameter controlling the trade-off between exploration and exploitation (*β*=2).

In each round, K-Means algorithm was applied to group the top-ranked SANPs (*γ*=0.1) into 20 clusters. From each cluster, the top 3 samples based on were selected, yielding a total of 60 candidates (**Supplementary Table S8**). These samples were then experimentally tested for antibacterial activity and hemolysis ratio. After several iterations of this process, the entire library was predicted and ranked using the final model. The top 60 candidates, based on their predicted scores *s*(*x*), were synthesized and characterized again to validate the model’s predictive performance and assess the effectiveness of the active learning approach.

### Minimum Inhibitory Concentration (MIC) assay

MRSA was incubated in TSB overnight (120 rpm, 37°C). The bacterial suspension was then diluted and calibrated to a concentration of 10^5^ CFU/mL. Purified SANPs were dissolved in DMSO to a concentration of 2.56 mg/mL. 195 μL MRSA suspension was added to the 96-well plate, and 5 μL tested SANPs solution was added to each well. The SANPs concentration was serially diluted two-fold, ranging from 32 μg/mL to 4 μg/mL. The plate was incubated for 9 h. TSB solution served as the control group. The MIC was determined as the lowest concentration that resulted in a completely clear solution.

### Determination of Half-hemolytic Concentration (HC_50_)

A 4% red blood cell suspension was prepared as described previously. For the assay, 195 μL of this suspension was added to each well of a 96-well plate, followed by the addition of 5 μL of purified SANPs solution. The final SANPs concentrations tested were 8, 16, 32, 64, 128, 200, 256, 400, and 512 μg/mL, respectively. The negative control group was added with 5 μL PBS, and the positive control group was added with 5 μL 0.4% Triton X-100. The plates were incubated at 37°C for 1 h, then centrifuged at 4°C (1500 rpm, 10 min). After centrifugation, 100 μL supernatant was transferred into a new 96-well plate. OD values at 576 nm were measured using a microplate reader, and the hemolysis ratio for each SANPs concentration was calculated. HC_50_ was determined by fitting the hemolysis ratio-concentration curve to identify the SANPs concentration at which 50% hemolysis occurred.

### Mouse pneumonia model

All animal experiments were conducted in accordance with relevant regulations and approved by the Animal Experimental Ethical Inspection Committee at Dr. Can Biotechnology (Zhejiang) Co., Ltd. On day 0, healthy female ICR mice (18-20 g) were anesthetized with isoflurane. MRSA suspension (10^8^ CFU/mL, 60 μL) was administered intratracheally. Successfully infected mice were randomly divided into 6 groups. On day 1, polymers H1, H2, H3, and SM were administered via tail vein injection (5 mg/kg). PBS served as the control. On day 2, the mice were euthanized, their lungs were excised for morphological observation. The lung tissue was homogenized for further bacterial colony counts.

### Checkerboard Assay

A checkerboard assay was performed to evaluate the synergistic effects of penicillin G sodium (PG) and the identified polymers (H1-3) against MRSA. The MICs of H1-3 were first determined, followed by the checkerboard assay conducted in 96-well microtiter plates. PG was serially diluted along the vertical axis from 128 µg/mL to 8 µg/mL, while the polymers were diluted along the horizontal axis: H1 and H3 ranged from 16 µg/mL to 1 µg/mL, and H2 ranged from 32 µg/mL to 2 µg/mL. The final row and column contained wells where both the polymer and PG concentrations were 0 µg/mL as controls. The plates were incubated at 37°C for 9 hours, and all experiments were performed in triplicate to ensure reproducibility and reliability.

### Preparation and characterization of H2-PG nanocomplexes (NCs)

The optimal formulation ratio for H2-PG nanocomplexes (NCs) was determined based on checkerboard assays. A stock solution of PG was prepared at 10 mg/mL in deionized water, while the polymer stock solution was 10.24 mg/mL in DMSO. Prior to use, the polymer solution was diluted with deionized water. For NC preparation, PG was diluted to 1 mg/mL in deionized water, and the polymer solution was diluted to the optimal concentration. Equal volumes of the PG and polymer solutions were mixed, vortexed for 30 s, and incubated at room temperature for 5 min to allow nanocomplex formation.

The hydrodynamic diameter of the NCs was measured using dynamic light scattering (DLS) on a Malvern Zetasizer Advance Pro (Blue Label). The zeta potential was also determined using the same instrument. The morphology of the NCs was examined by transmission electron microscopy (TEM).

The encapsulation efficiency (EE) and drug loading capacity (LC) of the antibiotic-loaded NCs were quantified using UV-Vis spectroscopy. NCs were prepared as described above and subsequently subjected to ultrafiltration using a 3 kDa molecular weight cutoff (MWCO) membrane to separate free PG from the nanocomplexes. The filtrate was collected and diluted appropriately, and its absorbance spectrum was recorded in the wavelength range of 185-220 nm using a UV-Vis spectrophotometer. Due to interference from the mixture, a slight shift in the position of the maximum absorption peak was observed; thus, the peak exhibiting the highest absorbance within this range was used to determine the concentration of unencapsulated PG. The EE and LC were calculated according to the following equations:

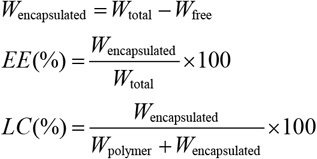

where *W*_total_ represents the initial total amount of PG, *W*_free_ is the amount of unencapsulated PG quantified from the filtrate, and *W*_polymer_ + *W*_encapsulated_ corresponds to the total mass of the nanocomplexes.

### Lethal MRSA Peritonitis Model

A lethal MRSA peritonitis model was established in 20 mice, randomly assigned to five groups (n = 4). Cyclophosphamide was administered intraperitoneally (150 mg/kg on day 1 and 100 mg/kg on day 3) to induce immunosuppression. On day 4, mice were intraperitoneally infected with 1 × 10^7^ CFU of MRSA. Post-infection, mice received intraperitoneal treatments: H2 (10 mg/kg), PG (10 mg/kg), SM (10 mg/kg), H2-PG nanocomplex (5 mg/kg), or PBS (100 μL, control). After 24 hours, mice were euthanized, and serum, spleen, lung, liver, and kidney samples were collected. Serum samples were used for the evaluation of inflammatory cytokine levels, while the harvested organs were analyzed for bacterial load and histopathological examination.

## Supporting information

Supplementary Information for the main text

## Data availability

The data that support the findings of this study are available within the main text and the Supplementary Information and can be obtained from the corresponding author upon request.

## Acknowledgements

This work was supported by the National Natural Science Foundation of China (52293381), Zhejiang Provincial Natural Science Foundation of China (LR25E030001) and the National Key Research and Development Program of China (2022YFB3807300). This work was also supported by Transvascular Implantation Devices Research Institute China (TIDRIC) under Grant No. KY012024007 and KY012024009.

## Contributions

P.Z. and J.J. conceptualized and supervised the project. Y.W., C.W., and X.S. designed the experiments, analyzed the data, and wrote the paper. Y.W. designed the overall framework and was responsible for the design, training, and analysis of the deep learning model. C. W. carried out the chemical synthesis and in vitro assays. X.S. investigated the self-assembly behavior of SANPs and their synergistic therapeutic effects with antibiotics. Y.C. and H.W. evaluated the performance of SANPs against clinically isolated multidrug-resistant bacteria. B.X. participated in the mouse peritonitis model experiments. Y.F.C. assisted with the experiments. W.D. and Y.H. contributed to the mouse pneumonia model experiments. L.Z. participated in experimental discussions. All authors reviewed and approved the final manuscript.

## Notes

### Competing Interest Statement

The authors have declared no competing interest.

