## Supplementary Information for the main text for "Deep learning guided discovery of antibacterial polymeric nanoparticles"

### Supplementary Methods

#### Measurement of Half Maximal Inhibitory Concentration ($IC_{50}$)

L929 mouse fibroblasts were cultured in DMEM supplemented with 10% serum. Cells were seeded into 96-well plates at a density of 8,000 cells per well. After 24 hours of incubation, the medium was removed, and 195 μL of fresh medium was added to each well, followed by 5 μL of pre-prepared SANPs solution at various concentrations. The cells were incubated for another 24 h. The medium was then discarded, and 100 μL of DMEM (containing 10% CCK-8 reagent) was added to each well. After 1 hour, OD values at 450 nm were measured with a microplate reader. The cell viability was calculated using the following formula:

$$\text{Cell viability}(\%) = \frac{OD_{\text{sample}} - OD_{\text{blank}}}{OD_{\text{negtive control}} - OD_{\text{blank control}}} \times 100$$

The cell viability-concentration curve was then fitted to determine the SANPs concentration at which 50% of the cells remained viable.

#### Bactericidal performance

The identified SANPs obtained via PolyCLOVER were designated as H1, H2, and H3. MRSA were cultured overnight in TSB at 37°C and diluted to $10^8$ CFU/mL. The MRSA suspension was centrifuged at 5000 rpm for 5 minutes, then the supernatant was removed. MRSA was resuspended in sterile PBS. H1, H2, and H3 were added to the suspension at a final concentration of 32 μg/mL for further 9 h incubation. After incubation, 100 μL of the mixture was sampled and spread onto Tryptic Soy Agar (TSA) plates to observe the formation of bacterial colonies. To quantify viable bacteria, 10 μL of the mixture was added to TSA plates and colonies were counted.

#### Bactericidal kinetics test

MRSA were cultured overnight in TSB at 37°C and diluted to $10^8$ CFU/mL. The suspension was centrifuged at 5000 rpm, and the supernatant was discarded. Then MRSA was resuspended in sterile PBS to a concentration of $10^5$ CFU/mL. H1, H2, H3, along with streptomycin (SM), were added to the MRSA suspension at a final concentration of 32 μg/mL. At designated time points (0, 4, 8, 10, 20, 30, 60, and 90 min), 10 μL of the solution was pipetted on TSA, and viable colonies were counted. DMSO was used as the control. The bacterial viability was calculated as the ratio of CFU count at each time point to the initial CFU count.

#### Antibacterial-resistance test

The antibacterial resistance assay followed a protocol similar to the MIC test. On day 1, the MIC was determined as described previously. Samples from turbid cultures at sub-MIC concentrations were collected, diluted 400-fold in TSB, and incubated overnight. The following day, the culture was further diluted 2000-fold in TSB and used for a new MIC determination. This cycle was repeated every 24 hours for 28 consecutive days.

#### Live/dead bacteria staining experiment

Staining experiments were performed using a live/dead bacteria assay kit (BBcellProbe®N01/PI, BestBio). 1 mL MRSA bacterial suspension (calibrated to $4 \times 10^8$ CFU/mL in sterile PBS) was treated with H1, H2, H3, and streptomycin (SM), at a final concentration of 64 μg/mL. After 3 minutes of co-incubation, the bacterial suspension was centrifuged, and the supernatant was discarded. The bacteria were then resuspended in the dye solution and incubated at room temperature for 15 minutes. Following incubation, the dye solution was removed by centrifugation, and the bacteria were washed with PBS and centrifuged again. The stained bacteria were observed with a laser confocal scanning microscope (Zeiss LSM 900).

**Membrane depolarization test**

2 mL fresh MRSA suspension was centrifuged and washed with HEPES buffer (5 mM, pH 7.4, containing 20 mM glucose). The supernatant was discarded, and the bacteria were resuspended and diluted to $10^7$ CFU/mL with HEPES buffer. The MRSA suspension was then treated with 0.4 μM 3,3'-dipropylthiadicarbocyanine iodide [$DiSC_3(5)$, Bidepharm] for 1 hour. Potassium chloride was added to the mixture to a final concentration of 0.1 M. 195 μL suspension was transferred into a 96-well plate. Fluorescence intensity (excitation at 622 nm, emission at 670 nm) was recorded using a microplate reader. Once the intensity value stabilized, 5 μL of H1, H2, H3, SM solution was immediately added to the suspension (final concentration at 64 μg/mL). The change in fluorescence intensity was then recorded. HEPES buffer served as the negative control, and Triton X-100 (final concentration 2.5%) was used as the positive control.

**TEM observation**

MRSA were resuspended in PBS and the suspension was incubated with H1, H2, or H3 at a final concentration of 64 μg/mL for 10 minutes. The suspension was subsequently centrifuged and washed with PBS to remove the supernatant. The MRSA were fixed overnight at 4°C with 2.5% glutaraldehyde solution. The fixed bacteria were washed three times with cold PBS, followed by incubation with 1% osmium tetroxide solution. After washing with cold PBS to remove residual osmium tetroxide, the bacteria were dehydrated through a series of ethanol gradients (30%, 50%, 70%, 90%, and 100%). The sample was then treated with acetone, embedded in resin, and sectioned into ultra-thin slices. The sections were observed using transmission electron microscopy (Hitachi HT7700).

**Mouse pneumonia model**

**SANPs distribution in mice.** H1, H2, and H3 were redissolved in DMF at 40 mg/mL. To 200 μL of each SANPs solution, Rhodamine B isothiocyanate (2 mg/mL, Sigma) was added, achieving a molar ratio of Rhodamine B isothiocyanate to identified SANPs of 2:1. Additionally, 5 μL of triethylamine was included. The mixture was stirred continuously overnight. The solution was then precipitated sequentially with ether/cyclohexane (1:1 vol%), ether/cyclohexane (4:1 vol%), and pure ether, yielding the Rhodamine B-labeled identified SANPs.

Pneumonia models were established in mice with the previously described method. On the first day, Rhodamine B-labeled H1, H2, and H3 were administered via tail vein injection (5 mg/kg), with PBS as the control. Two hours post-injection, mice were euthanized, and major organs were collected. Fluorescence imaging was performed using a small animal fluorescence imager (PerkinElmer).

**Tissue biocompatibility research.** Healthy female ICR mice were injected via the tail vein with H1, H2 and H3 (5 mg/kg), with PBS serving as the control. 24 h post-injection, the mice were euthanized. The heart, liver, spleen, lung, and kidney were excised and subjected to H&E staining.

Tissue morphology was examined to assess any potential toxicity of the SANPs to the mice.

**Lethal MRSA Peritonitis Model**

**Hematoxylin and eosin (H&E) staining.** Following euthanasia, spleen, liver, lung, and kidney tissues were harvested and fixed in 4% paraformaldehyde overnight. The fixed tissues were paraffin-embedded and sectioned at a thickness of 5 μm. The sections were then deparaffinized in xylene and rehydrated through a graded ethanol series. For histological assessment, the sections were stained with hematoxylin and eosin (H&E) to differentiate nuclear and cytoplasmic structures. Images were acquired using a light microscope, with three randomly selected fields analyzed per section.

**Analysis of bacterial load.** Tissue samples were washed with phosphate-buffered saline (PBS), homogenized, and subjected to low-speed centrifugation at 100 rpm, after which the supernatant was collected for bacterial quantification. The samples were serially diluted tenfold, and 100 μL of each dilution was plated on nutrient agar for CFU quantification. CFUs were enumerated and expressed as CFU per gram of tissue.

**Elisa.** Mouse blood was centrifuged at 4,000 g for 15 minutes at 4°C, and serum was collected and frozen at -80°C. The serum was analyzed by using ELISA kits for mouse IL-6, TNF-α according to the manufacturer's guidelines.

**Supplementary Figs**

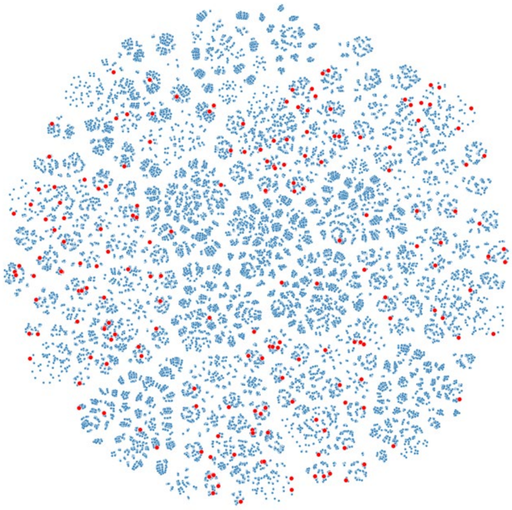

**Fig. S1 | t-SNE visualization of the combinatorial library.** Each point represents a nanoparticle candidate, and red points indicate those selected in the initial sampling.

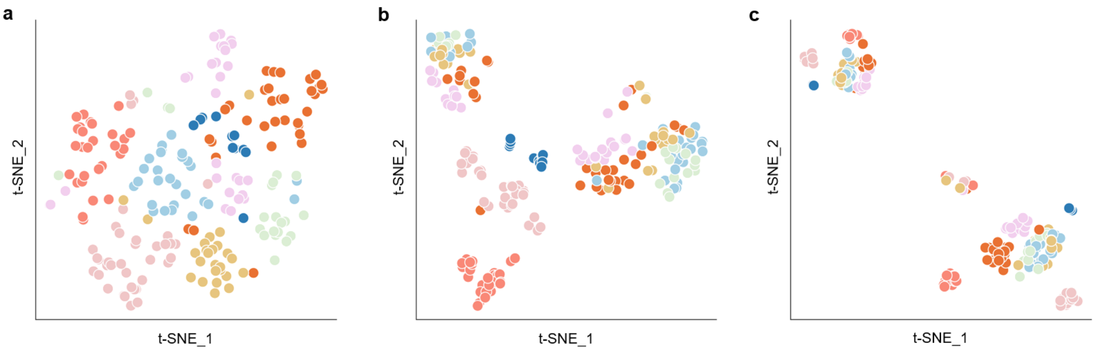


**Fig. S2 | t-SNE visualization of representations from pretrained models, colored by diacrylate type. a**, Two-stage pre-trained model. **b**, Single-stage pretrained model. The model is not pre-trained on the constructed combinatorial library. **c**, Un-pretrained model.

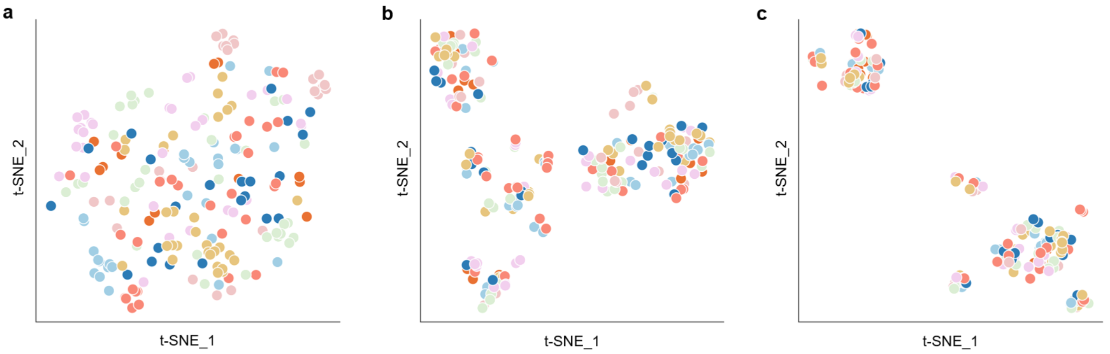


**Fig. S3 | t-SNE visualization of representations from pretrained models, colored by positive charged amine type. a**, Two-stage pre-trained model. **b**, Single-stage pretrained model. The model is not pre-trained on the constructed combinatorial library. **c**, Un-pretrained model.

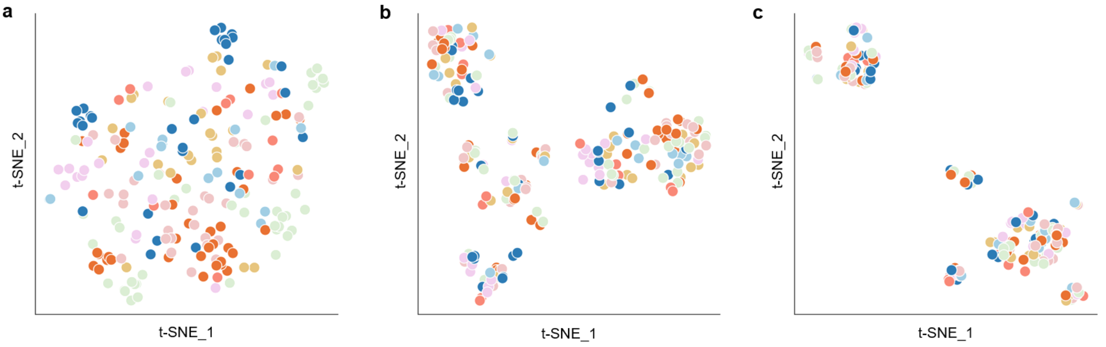


**Fig. S4 | t-SNE visualization of representations from pretrained models, colored by hydrophobic amine type. a**, Two-stage pre-trained model. **b**, Single-stage pretrained model. The model is not pre-trained on the constructed combinatorial library. **c**, Un-pretrained model.

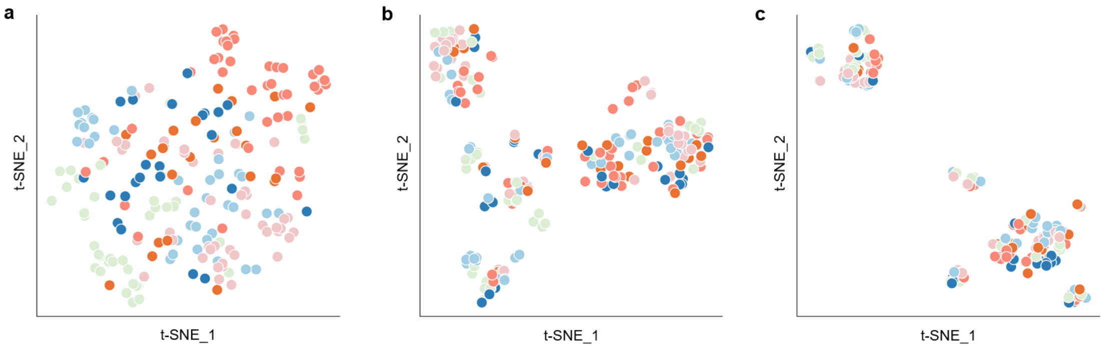


**Fig. S5 | t-SNE visualization of representations from pretrained models, colored by hydrophilic amine type. a**, Two-stage pre-trained encoder. **b**, Single-stage pretrained encoder. The model is not pre-trained on the constructed combinatorial library. **c**, Un-pretrained encoder.

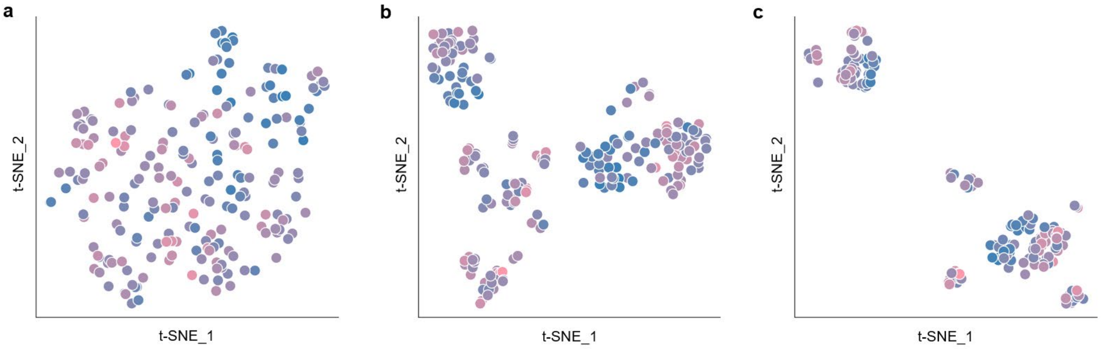


**Fig. S6 | t-SNE visualization of representations from pretrained models, colored by average LogP. a**, Two-stage pre-trained model. **b**, Single-stage pretrained model. The model is not pre-trained on the constructed combinatorial library. **c**, Un-pretrained model.

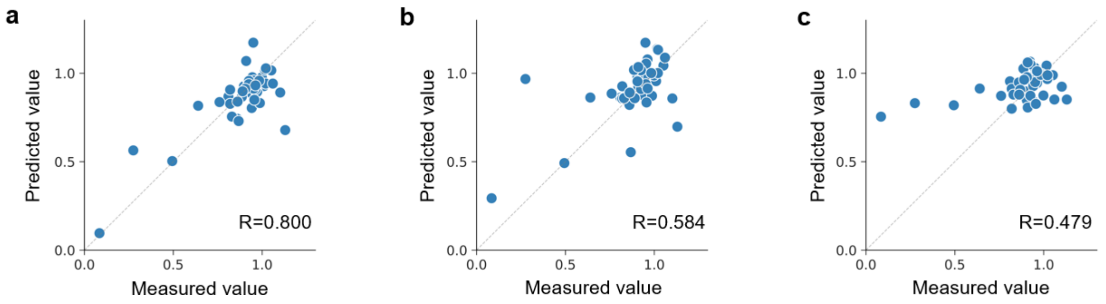


**Fig. S7 | The measured antibacterial property versus predicted plots for each model. a**, The pre-trained model. **b**, The weighted sum model. **c**, The un-pretrained model. The dashed line represents ideal case y=x. The Pearson correlation coefficient is shown in the lower right corner.

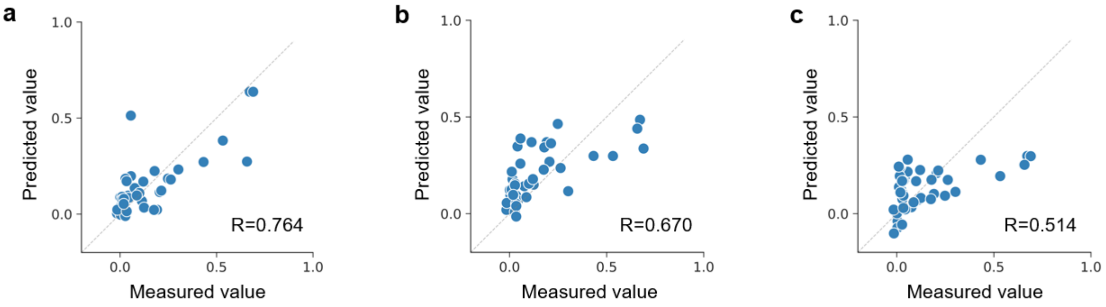


**Fig. S8 | The measured hemolytic property versus predicted plots for each model. a**, The pre-trained model. **b**, The weighted sum model. **c**, The un-pretrained model. The dashed line represents ideal case y=x. The Pearson correlation coefficient is shown in the lower right corner.

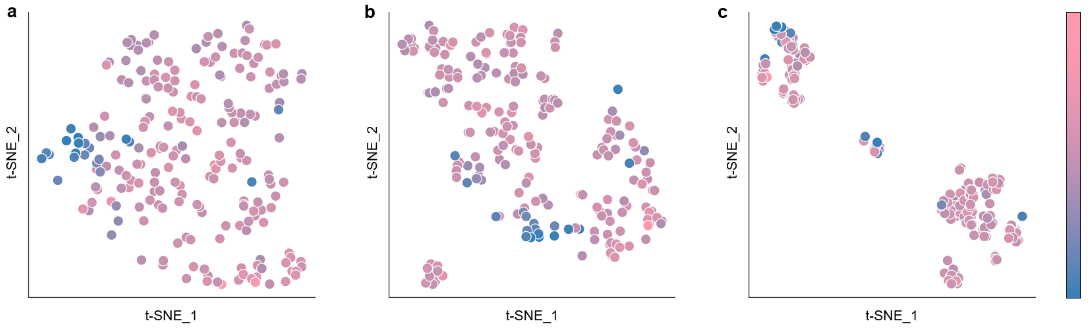


**Fig. S9 | t-SNE visualization of representations from fine-tuned models, colored by antibacterial property.** **a**, Fine-tuned two-stage pretrained model. **b**, Fine-tuned single-stage pretrained model. **c**, Fine-tuned model without any pre-training.

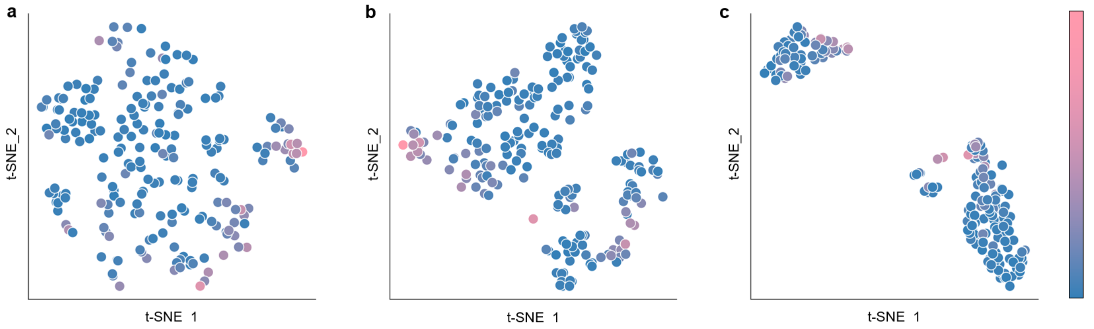


**Fig. S10 | t-SNE visualization of representations from fine-tuned models, colored by hemolytic property.** **a**, Fine-tuned two-stage pretrained model. **b**, Fine-tuned single-stage pretrained model. **c**, Fine-tuned model without any pre-training.

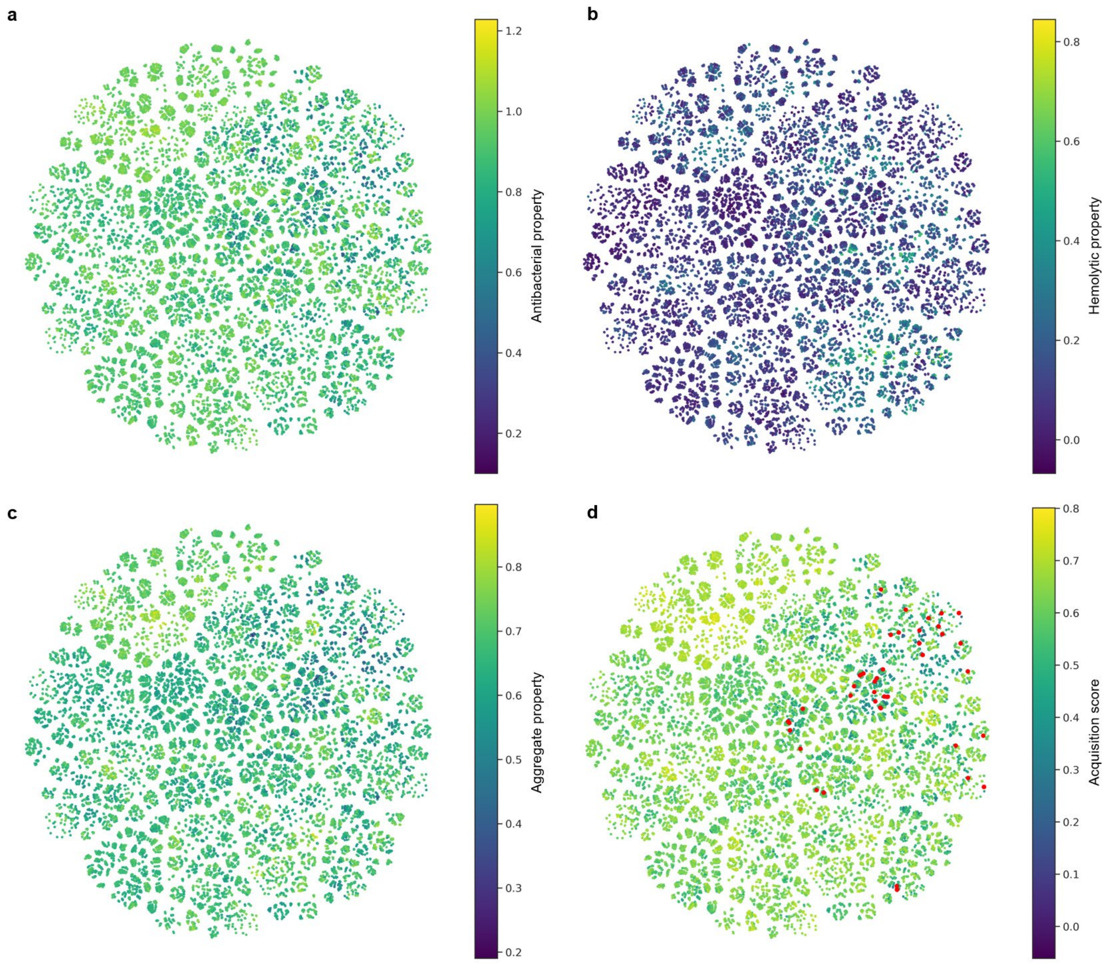


**Fig. S11 | t-SNE visualization of the library in Round 1. a**, Predicted antibacterial activity of unlabeled samples in the constructed library. **b**, Predicted hemolytic property of unlabeled samples in the constructed library. **c**, Aggregated score obtained by weighted summation of the two predicted properties. **d**, Acquisition score used for sample selection. Red points indicate the candidates recommended by PolyCLOVER for labeling in the next round.

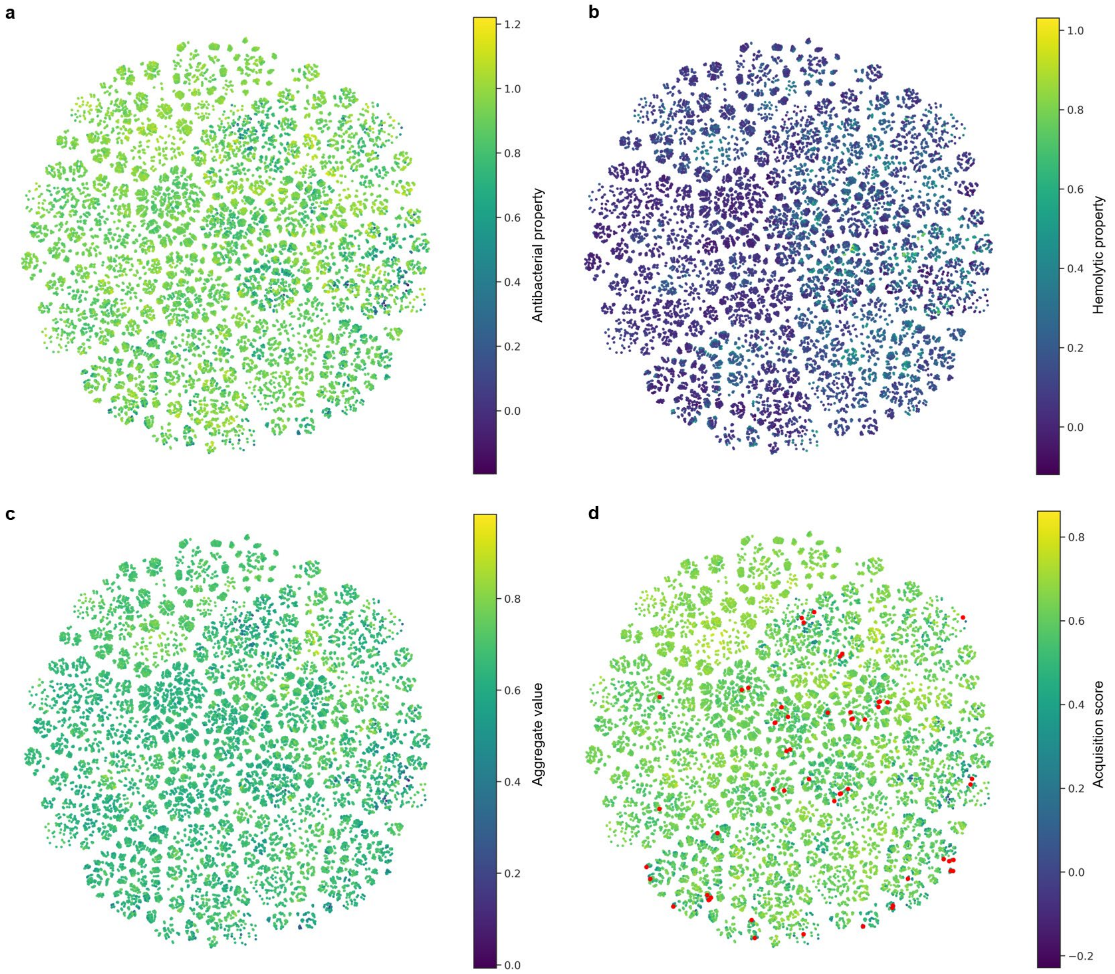


**Fig. S12 | t-SNE visualization of the library in Round 2. a**, Predicted antibacterial activity of unlabeled samples in the constructed library. **b**, Predicted hemolytic property of unlabeled samples in the constructed library. **c**, Aggregated score obtained by weighted summation of the two predicted properties. **d**, Acquisition score used for sample selection. Red points indicate the candidates recommended by PolyCLOVER for labeling in the next round.

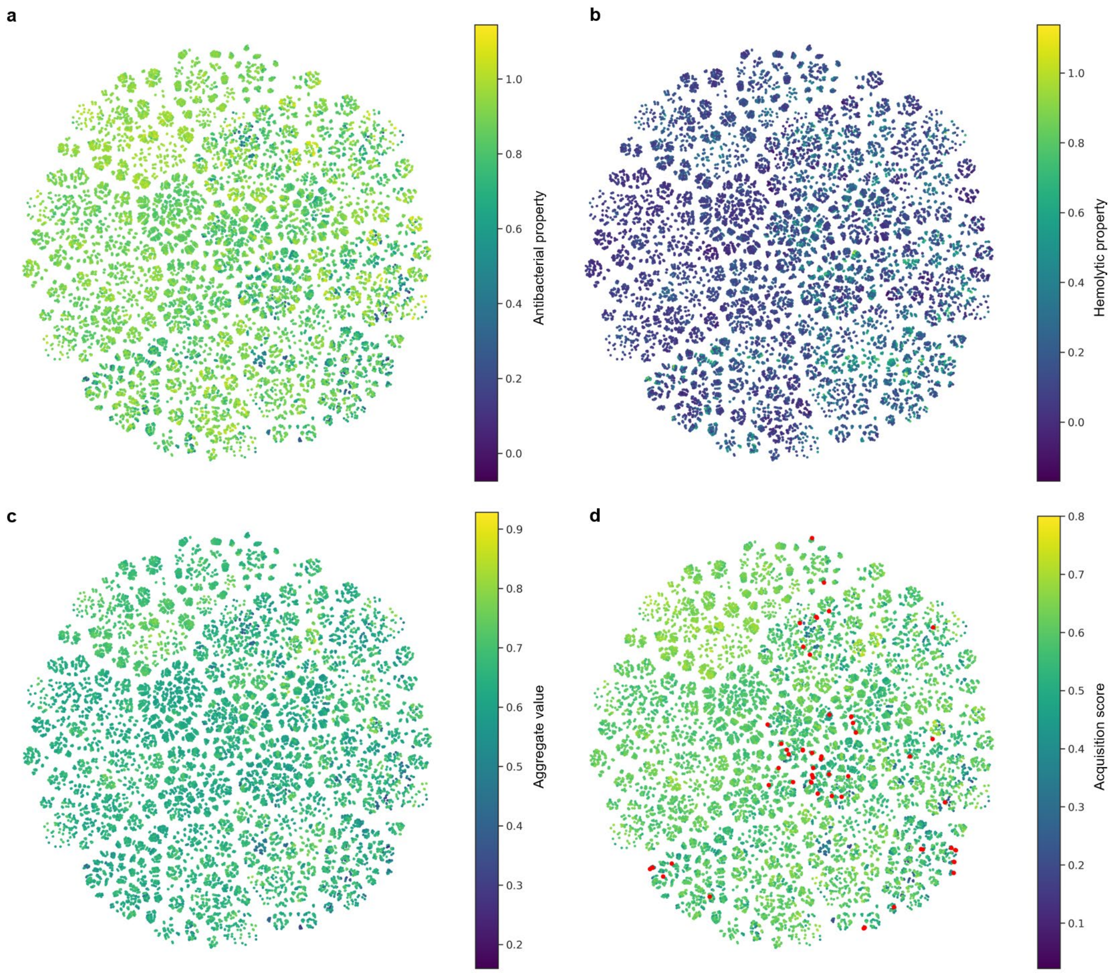


**Fig. S13 | t-SNE visualization of the library in Round 3. a**, Predicted antibacterial activity of unlabeled samples in the constructed library. **b**, Predicted hemolytic property of unlabeled samples in the constructed library. **c**, Aggregated score obtained by weighted summation of the two predicted properties. **d**, Acquisition score used for sample selection. Red points indicate the candidates recommended by PolyCLOVER for labeling in the next round.

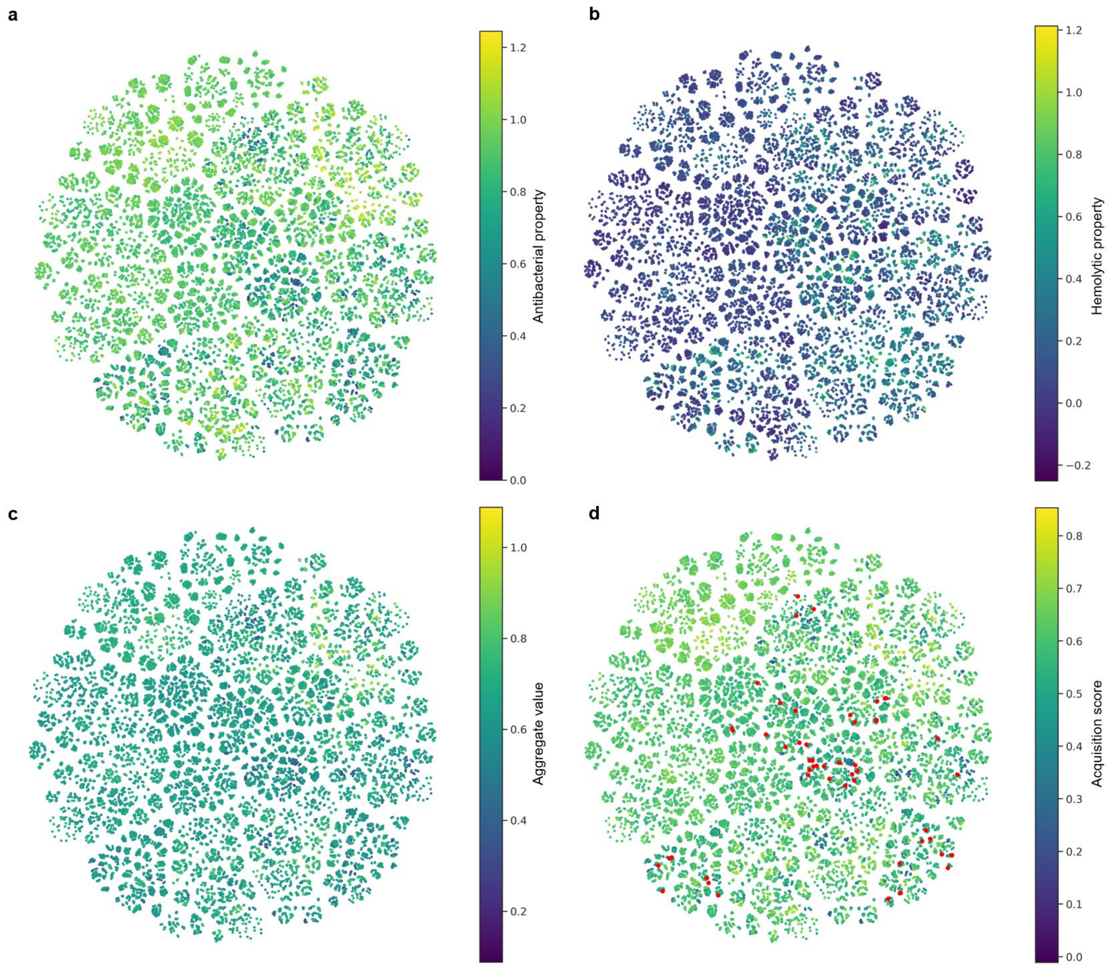


**Fig. S14 | t-SNE visualization of the library in Round 4. a**, Predicted antibacterial activity of unlabeled samples in the constructed library. **b**, Predicted hemolytic property of unlabeled samples in the constructed library. **c**, Aggregated score obtained by weighted summation of the two predicted properties. **d**, Acquisition score used for sample selection. Red points indicate the candidates recommended by PolyCLOVER for labeling in the next round.

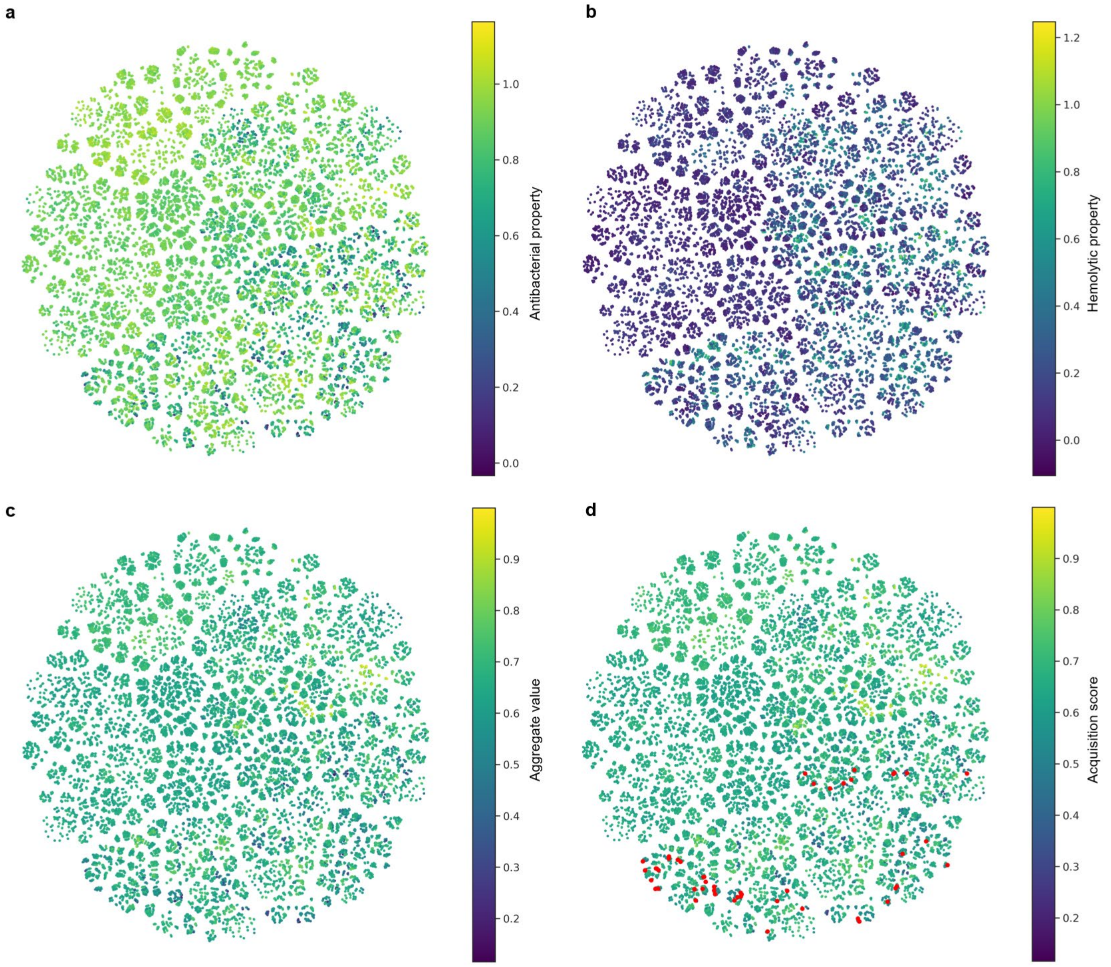


**Fig. S15 | t-SNE visualization of the library in the final round. a**, Predicted antibacterial activity of unlabeled samples in the constructed library. **b**, Predicted hemolytic property of unlabeled samples in the constructed library. **c**, Aggregated score obtained by weighted summation of the two predicted properties. **d**, Acquisition score used for sample selection. Red points indicate the candidates recommended by PolyCLOVER for final labeling.

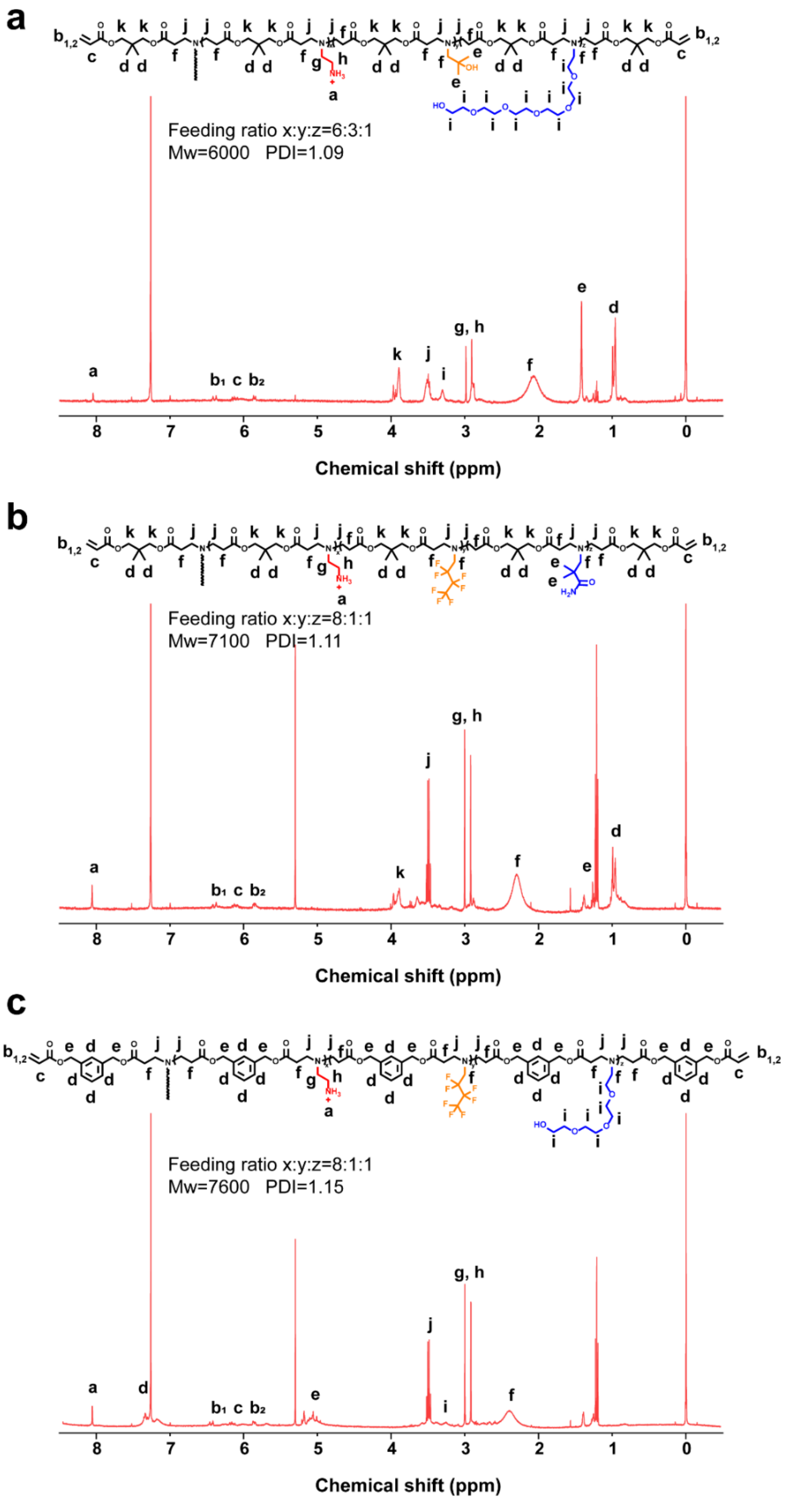


**Fig. S16 | Structures and nuclear magnetic resonance spectra of the identified SANPs (a) H1, (b) H2 and (c) H3.**

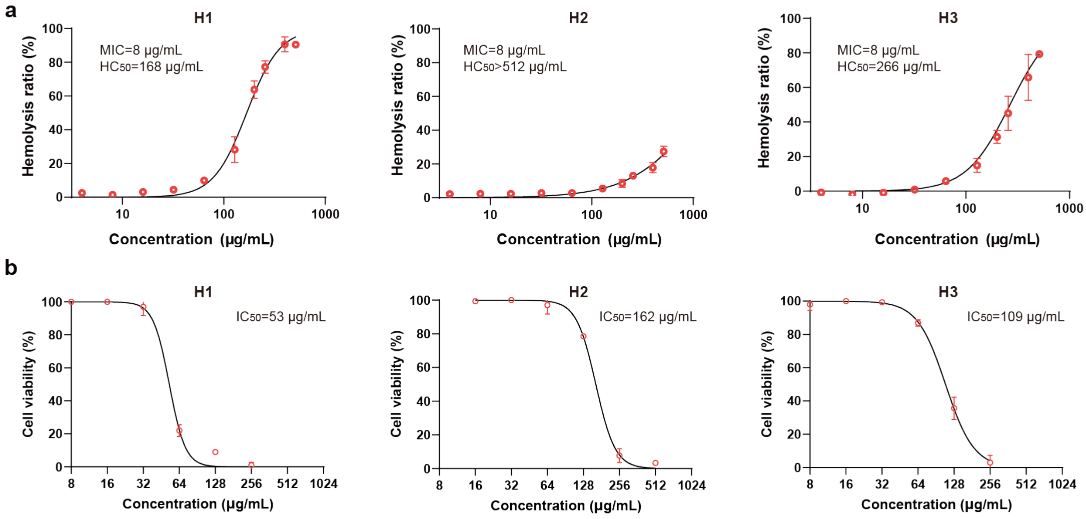


**Fig. S17 | (a) Hemolysis ratio-concentration curves and (b) cell viability-concentration curves of the identified SANPs.** Data are shown as mean±s.d. (n = 3 independent replicates).

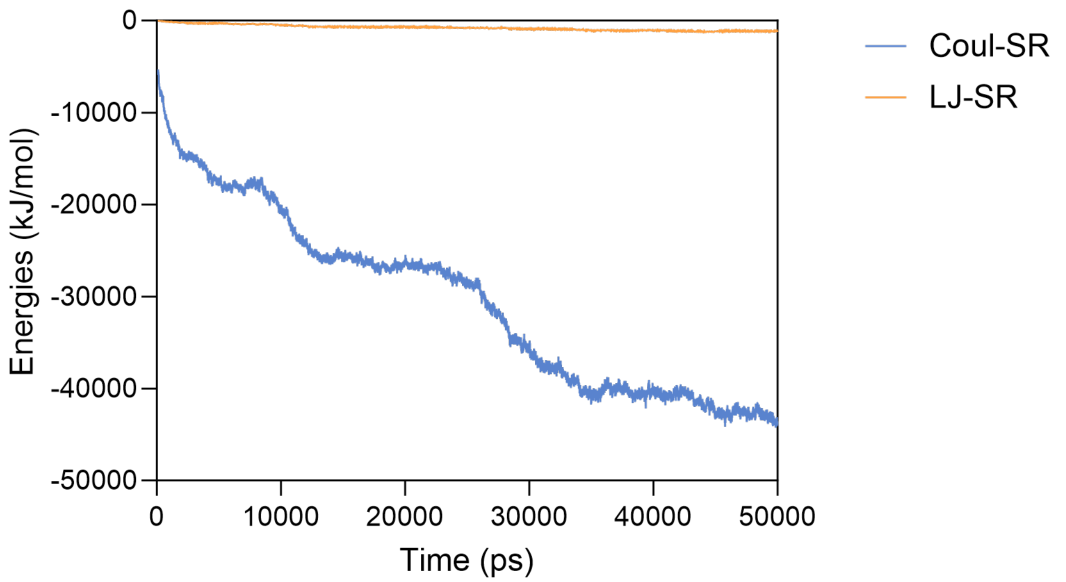


**Fig. S18 | Time-dependent changes in the interactions between H2 and the cell membrane during the assembly process observed in molecular dynamics simulations.** The time evolution of electrostatic interactions (Coul-SR) and van der Waals interactions (LJ-SR) between H2 and the cell membrane is shown. Positive values represent repulsive forces, while negative values indicate attractive interactions. Both types of interactions display an overall attractive trend. Notably, van der Waals interactions remain relatively stable with minor fluctuations, whereas electrostatic interactions become increasingly negative over time, suggesting a progressively stronger electrostatic attraction between H2 and the membrane.

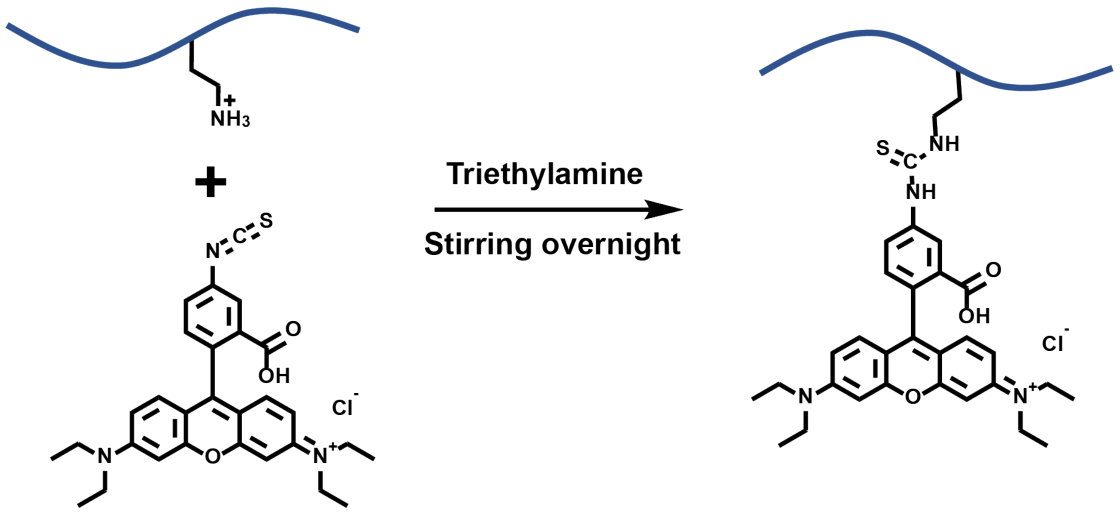


**Fig. S19 | Reaction to graft Rhodamine B isothiocyanate to the identified SANPs.**

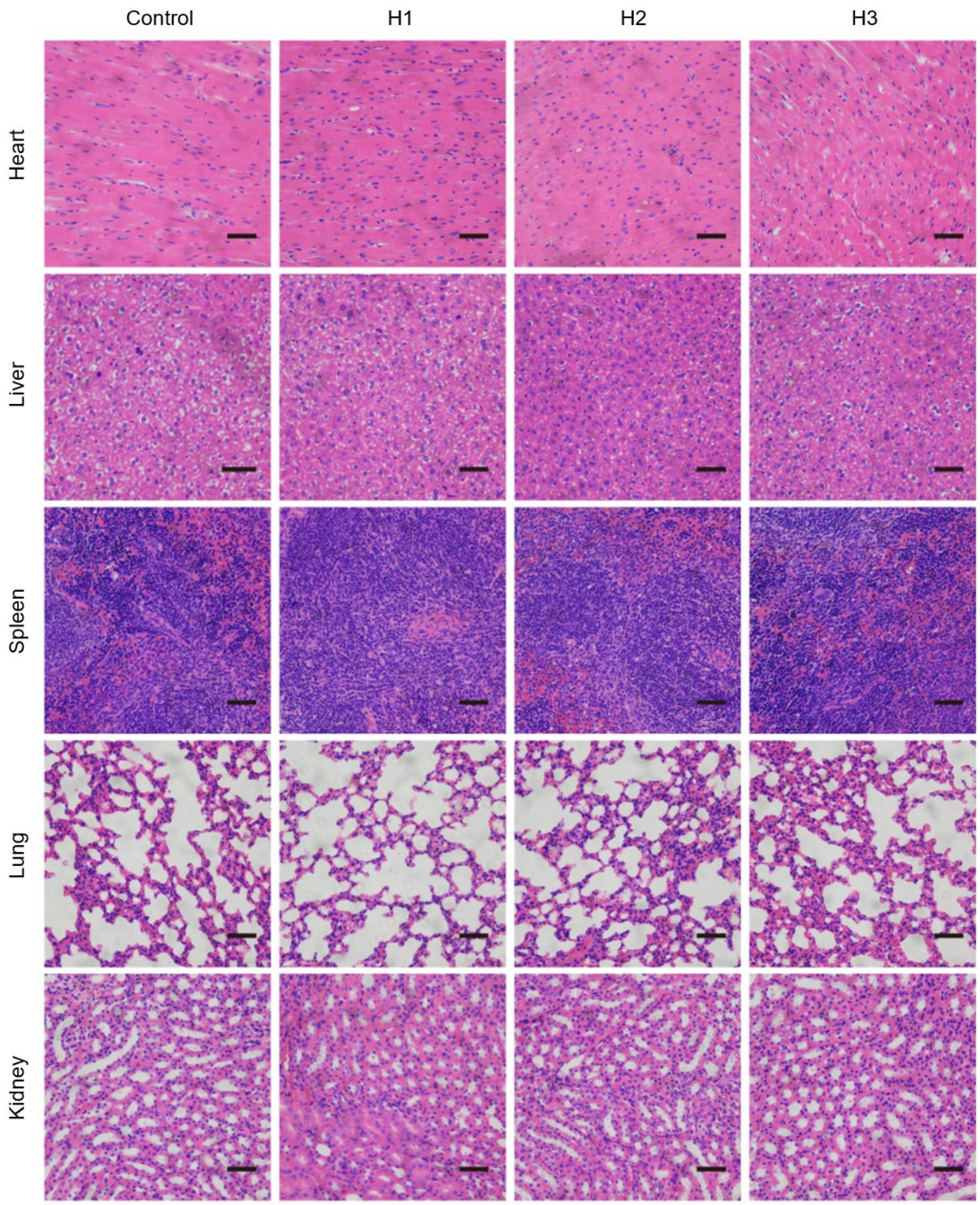


**Fig. S20 | H&E staining of mice organs (heart, liver, spleen, lung, kidney).** Scale bar: 50 μm.

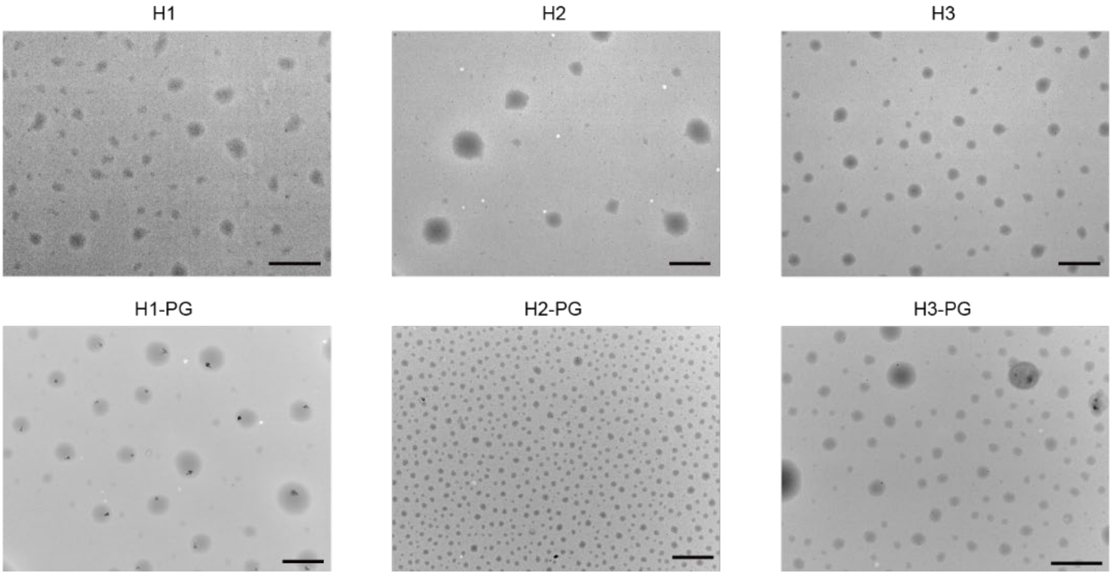


**Fig. S21 | TEM images of identified SANPs before and after PG loading.** TEM images of identified SANPs before (top) and after (bottom) loading with PG. Upon PG loading, H2 became significantly smaller and more uniform in size. Experiments were performed in triplicate with consistent results; representative images are shown. Scale bar: 1 μm.

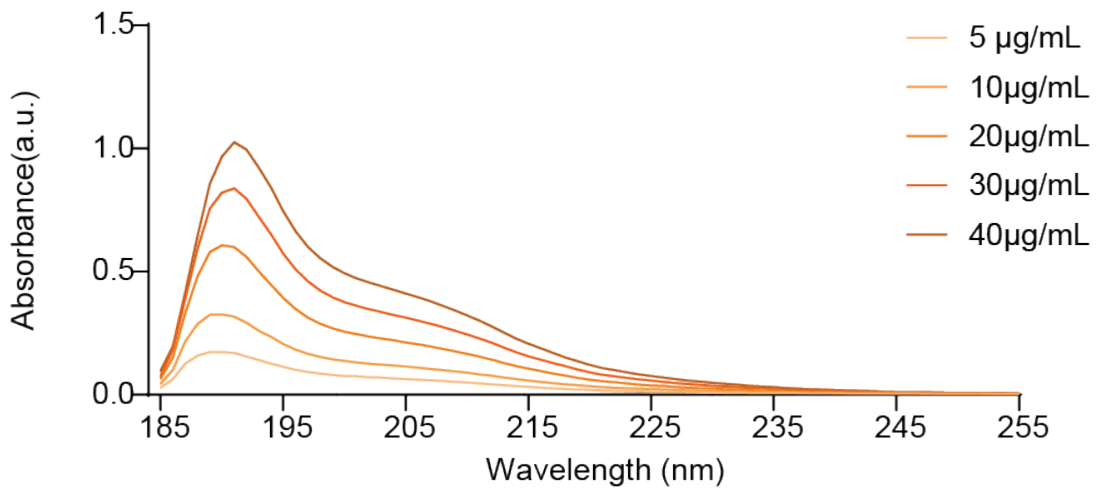


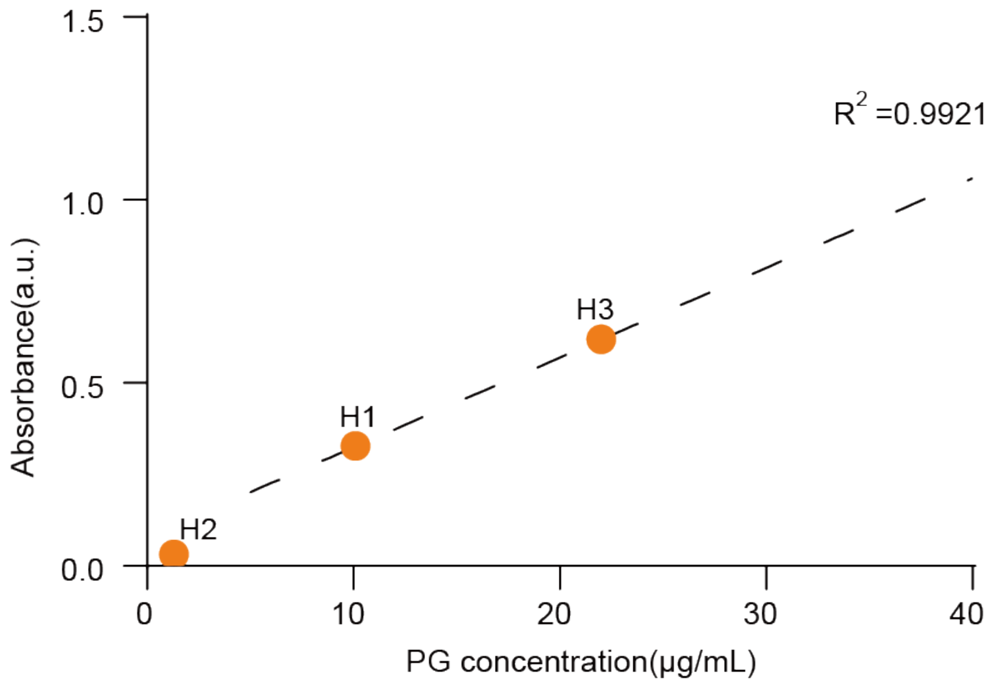


**Fig. S22 | Measurement of the encapsulation efficiency and loading capacity of PG in identified SANPs using UV-Vis spectroscopy.** A standard curve is constructed using PG standards at different concentrations. The ultrafiltrate of the mixture of identified SANPs and PG is used as the sample for quantifying the amount of unencapsulated PG.

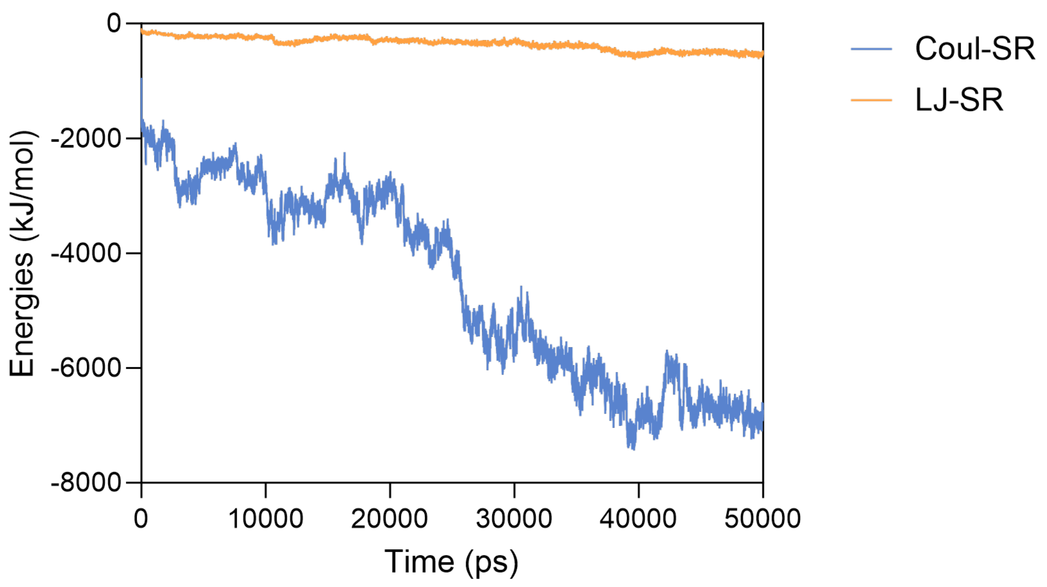


**Fig. S23 | Time-dependent changes in the interactions between five penicillin molecules and five H2 molecules during the assembly process in molecular dynamics simulations.** Time evolution of the electrostatic interactions (Coul-SR) and van der Waals interactions (LJ-SR) between H2 and PG. Positive values indicate repulsion between the two molecules, while negative values indicate attraction. Both interactions exhibit an attractive trend, with van der Waals interactions showing slight variation and electrostatic interactions gradually increasing, indicating an increasingly stronger attraction.

**Supplementary Tables**

**Table S1 | Reagents for the library construction**

| Serial number | Name | CAS | SMILES | Producer |
|---|---|---|---|---|
| A1 | Ethylene glycol diacrylate | 2274-11-5 | C=CC(OCCOC(C=C)=O)=O | Bidepharm |
| A2 | Neopentyl glycol diacrylate | 2223-82-7 | CC(C)(COC(C=C)=O)COC(C=C)=O | Bidepharm |
| A3 | 1,4-Butanediol diacrylate | 1070-70-8 | C=CC(OCCCCOC(C=C)=O)=O | Bidepharm |
| A4 | 1,5-Pentanediol Diacrylate | 36840-85-4 | C=CC(OCCCCCOC(C=C)=O)=O | Bidepharm |
| A5 | 1,6-Hexanediol Diacrylate | 13048-33-4 | C=CC(OCCCCCCOC(C=C)=O)=O | D&B |
| A6 | Triethylene glycol diacrylate | 1680-21-3 | C=CC(OCCOCCOCCOC(C=C)=O)=O | Bidepharm |
| A7 | Tripropylene Glycol Diacrylate | 42978-66-5 | CC(OCC(OCC(OC(C=C)=O)C)C)COC(C=C)=O | Bidepharm |
| A8 | 1,3-Phenylenebis(methylene) diacrylate | 22757-16-0 | C=CC(OCC1=CC=CC(COC(C=C)=O)=C1)=O | Bidepharm |
| B1 | N,N-Dimethylethylenediamine | 108-00-9 | NCCN(C)C | Aladdin |
| B2 | 3-Diethylaminopropylamine | 104-78-9 | NCCCN(CC)CC | Aladdin |
| B3 | 3-(Dibutylamino)propylamine | 102-83-0 | NCCCN(CCCC)CCCC | Bidepharm |
| B4 | 1-(2-Aminoethyl)pyrrolidine | 7154-73-6 | NCCN1CCCC1 | Aladdin |
| B5 | N-(2-Aminoethyl)piperidine | 27578-60-5 | NCCN1CCCCC1 | Bidepharm |
| B6 | 1-(3-Aminopropyl)-4-methylpiperazine | 4572-03-6 | CN1CCN(CCCN)CC1 | Bidepharm |
| B7 | 4-Pyridinemethaneamine | 3731-53-1 | NCC1=CC=NC=C1 | Bidepharm |
| B8 | 3-(2-Aminoethyl)pyridi | 20173-24-4 | NCCC1=CC=CN=C1 | Bidepharm |

| | | | | |
|---|---|---|---|---|
| | ne | | | |
| B9 | 4-(2-Aminoethyl)morpholine | 2038-03-1 | NCCN1CCOCC1 | Bidepharm |
| B10 | N-Boc-ethylenediamine | 57260-73-8 | NCCNC(OC(C)(C)C)=O | Bidepharm |
| B11 | N-Boc-2,2-dimethyl-1,3-diaminopropane | 292606-35-0 | NCC(C)(C)CNC(OC(C)(C)C)=O | Bidepharm |
| B12 | N-(5-Aminoamyl)Carbamic Acid Tert-Butyl Ester | 51644-96-3 | O=C(OC(C)(C)C)NCCCCCN | Aladdin |
| B13 | tert-Butyl (8-aminooctyl)carbamate | 88829-82-7 | O=C(OC(C)(C)C)NCCCCCCCCN | Bidepharm |
| B14 | N-(tert-Butoxycarbonyl)-2,2'-(ethylenedioxy)diethylamine | 153086-78-3 | NCCOCCOCCNC(OC(C)(C)C)=O | Bidepharm |
| B15 | tert-Butyl (14-amino-3,6,9,12-tetraoxatetradecyl)carbamate | 811442-84-9 | O=C(OC(C)(C)C)NCCOCCOCCOCCOCCN | Bidepharm |
| B16 | tert-Butyl N-[3-(Aminomethyl)benzyl]carbamate | 108467-99-8 | O=C(OC(C)(C)C)NCC1=CC=CC(CN)=C1 | Bidepharm |
| C1 | 1-hexanamine | 111-26-2 | CCCCCCN | Macklin |
| C2 | Octylamine | 111-86-4 | NCCCCCCCC | Leyan |
| C3 | 1-Aminodecane | 2016-57-1 | CCCCCCCCCCN | Bidepharm |
| C4 | Cyclopentylmethanamine | 6053-81-2 | NCC1CCCC1 | Bidepharm |
| C5 | 2-Cyclohexylethylamine | 4442-85-7 | NCCC1CCCCC1 | Bidepharm |
| C6 | Benzylamine | 100-46-9 | NCC1=CC=CC=C1 | Energy Chemical |
| C7 | 3-Phenyl-1-propylamine | 2038-57-5 | NCCCC1=CC=CC=C1 | Energy Chemical |
| C8 | 4-Phenylbutylamine | 13214-66-9 | NCCCCC1=CC=CC=C1 | Macklin |
| C9 | Isoamylamine | 107-85-7 | NCCC(C)C | Macklin |
| C10 | 2-Ethylhexylamine | 104-75-6 | NCC(CC)CCCC | Aladdin |

| | | | | |
|---|---|---|---|---|
| C11 | 3,3-Diethoxypropylamine | 41365-75-7 | NCCC(OCC)OCC | Bidepharm |
| C12 | 4-Methoxybenzylamine | 2393-23-9 | NCC1=CC=C(OC)C=C1 | Aladdin |
| C13 | 1-Amino-2-methylpropan-2-ol | 2854-16-2 | CC(O)(C)CN | Bidepharm |
| C14 | 3-Methoxypropylamine | 5332-73-0 | NCCCOC | Bidepharm |
| C15 | 5-Aminopentan-1-ol | 2508-29-4 | OCCCCCN | Macklin |
| C16 | 2,2,3,3,4,4,4-Heptafluorobutylamine | 374-99-2 | NCC(F)(F)C(F)(F)C(F)(F)F | Bidepharm |
| D1 | 2-(2-(2-Aminoethoxy)ethoxy)ethanol | 6338-55-2 | NCCOCCOCCO | Energy Chemical |
| D2 | 1-Amino-3,6,9-trioxaundecanyl-11-ol | 86770-74-3 | NCCOCCOCCOCCO | Bidepharm |
| D3 | 17-Amino-3,6,9,12,15-pentaoxaheptadecanol | 39160-70-8 | NCCOCCOCCOCCOCCOCCO | Bidepharm |
| D4 | 3-Amino-1,2-propanediol | 616-30-8 | OCC(O)CN | Bidepharm |
| D5 | 3-Amino-2,2-dimethylpropanamide | 324763-51-1 | O=C(N)C(C)(C)CN | Bidepharm |
| D6 | 2-aminopropanamide | 4726-84-5 | CC(N)C(N)=O | Bidepharm |

**Table S2 | Feeding ratio of poly (β-amino ester) combinatorial library.**

| Index | Diacrylate | Amine A | Amine B | Amine C |
|---|---|---|---|---|
| 1 | 12 | 2 | 2 | 6 |
| 2 | 12 | 2 | 4 | 4 |
| 3 | 12 | 2 | 6 | 2 |
| 4 | 12 | 4 | 2 | 4 |
| 5 | 12 | 4 | 4 | 2 |
| 6 | 12 | 6 | 1 | 3 |
| 7 | 12 | 6 | 3 | 1 |
| 8 | 12 | 8 | 1 | 1 |

**Table S3 | Test performance of different backbones on antibacterial and hemolytic tasks.**

| **Backbone** | **RMSE of antibacterial property** | **RMSE of hemolytic property** | **Average RMSE** |
|---|---|---|---|
| GAT | 0.1837±0.0070 | 0.1993±0.0111 | 0.1915 |
| GIN | 0.1887±0.0035 | 0.1897±0.0012 | 0.1892 |
| AttentiveFP | 0.1903±0.0021 | 0.1867±0.0023 | 0.1885 |
| MPNN | 0.1913±0.0012 | 0.1847±0.0006 | 0.1880 |
| Weave | 0.1933±0.0055 | 0.1823±0.0012 | 0.1878 |
| Gated GCN | 0.1970±0.0026 | 0.1767±0.0035 | 0.1868 |
| RF | 0.1861±0.0037 | 0.1818±0.0053 | 0.1839 |
| GAT v2 | 0.1877±0.0025 | 0.1787±0.0023 | 0.1832 |
| GCN | 0.1817±0.0038 | 0.1797±0.0040 | 0.1807 |
| GraphGPS | **0.1730±0.0035** | 0.1740±0.0026 | 0.1735 |
| LiGhT | 0.1773±0.0051 | **0.1587±0.0150** | **0.1680** |

The mean and standard deviation of the test root mean square error on three independent runs are reported. The best performance for each task is shown in bold.

**Table S4 | Structures and detailed activities of SANPs meeting preliminary criteria.**

| Diacrylate (A) | Positive charged amine (B) | Hydrophobic amine (C) | Hydrophilic amine (D) | Ratio index | MIC (μg/mL) | Selectivity index |
|---|---|---|---|---|---|---|
| A3 | B5 | C3 | D3 | 5 | 32 | 9.8 |
| A4 | B5 | C3 | D4 | 5 | 32 | 5.1 |
| A5 | B5 | C3 | D5 | 5 | 32 | 9.1 |
| A2 | B10 | C2 | D6 | 6 | 32 | 6.8 |
| A2 | B10 | C10 | D5 | 8 | 16 | 15 |
| A2 | B10 | C10 | D6 | 8 | 16 | 14.9 |
| A2 | B10 | C16 | D3 | 7 | 32 | 14.1 |
| A2 | B10 | C16 | D6 | 8 | 32 | 15.5 |
| A3 | B10 | C2 | D5 | 8 | 32 | 16 |
| A4 | B10 | C2 | D6 | 8 | 32 | 6.4 |
| A4 | B10 | C10 | D5 | 8 | 32 | 12.7 |
| A5 | B10 | C2 | D5 | 8 | 16 | 8.3 |
| A5 | B10 | C9 | D5 | 8 | 16 | 16.9 |
| A5 | B10 | C13 | D3 | 8 | 32 | 7.6 |
| A5 | B10 | C13 | D5 | 6 | 32 | 8.7 |
| A5 | B10 | C13 | D5 | 8 | 32 | 4.5 |
| A5 | B10 | C16 | D5 | 6 | 32 | 16 |
| A5 | B10 | C16 | D5 | 8 | 16 | 17.9 |
| A8 | B10 | C10 | D3 | 8 | 32 | 7.5 |
| A8 | B10 | C10 | D5 | 8 | 32 | 8.7 |
| A8 | B10 | C16 | D3 | 8 | 32 | 16 |
| A8 | B10 | C16 | D5 | 6 | 32 | 16 |
| A8 | B10 | C16 | D5 | 8 | 32 | 13.1 |
| A2 | B10 | C1 | D5 | 8 | 16 | 18.8 |
| A2 | B10 | C1 | D6 | 8 | 32 | 8.5 |
| A2 | B10 | C2 | D3 | 8 | 16 | 13.2 |
| A2 | B10 | C2 | D5 | 7 | 16 | 8.6 |
| A2 | B10 | C2 | D5 | 8 | 32 | 9.9 |
| A2 | B10 | C2 | D6 | 7 | 16 | 4.3 |
| A2 | B10 | C3 | D3 | 8 | 16 | 12.6 |
| A2 | B10 | C3 | D5 | 5 | 16 | 4.2 |
| A2 | B10 | C3 | D5 | 7 | 16 | 5.4 |
| A2 | B10 | C3 | D5 | 8 | 16 | 8.7 |
| A2 | B10 | C3 | D6 | 5 | 32 | 4.2 |
| A2 | B10 | C3 | D6 | 7 | 16 | 6.7 |
| A2 | B10 | C3 | D6 | 8 | 16 | 7.9 |
| A2 | B10 | C8 | D5 | 8 | 8 | 16.6 |
| A2 | B10 | C8 | D6 | 8 | 16 | 12.4 |
| A2 | B10 | C10 | D3 | 8 | 16 | 17.5 |
| A2 | B10 | C10 | D5 | 6 | 64 | 1.7 |

| | | | | | | |
|---|---|---|---|---|---|---|
| A2 | B10 | C10 | D6 | 7 | 32 | 1.9 |
| A2 | B10 | C13 | D3 | 7 | 8 | 21 |
| A2 | B10 | C13 | D3 | 8 | 16 | 14.2 |
| A2 | B10 | C13 | D5 | 5 | 32 | 4.7 |
| A2 | B10 | C13 | D5 | 6 | 16 | 10.2 |
| A2 | B10 | C13 | D5 | 7 | 32 | 4.1 |
| A2 | B10 | C13 | D5 | 8 | 32 | 6.6 |
| A2 | B10 | C13 | D6 | 5 | 32 | 4.8 |
| A2 | B10 | C13 | D6 | 6 | 32 | 5.9 |
| A2 | B10 | C13 | D6 | 7 | 16 | 10.2 |
| A2 | B10 | C13 | D6 | 8 | 32 | 2.9 |
| A2 | B10 | C16 | D3 | 5 | 32 | 16 |
| A2 | B10 | C16 | D3 | 6 | 32 | 16 |
| A2 | B10 | C16 | D3 | 8 | 32 | 16 |
| A2 | B10 | C16 | D5 | 3 | 64 | 8 |
| A2 | B10 | C16 | D5 | 4 | 32 | 16 |
| A2 | B10 | C16 | D5 | 5 | 64 | 8 |
| A2 | B10 | C16 | D5 | 6 | 64 | 8 |
| A2 | B10 | C16 | D5 | 7 | 32 | 16 |
| A2 | B10 | C16 | D5 | 8 | 8 | 64 |
| A2 | B10 | C16 | D6 | 3 | 64 | 8 |
| A2 | B10 | C16 | D6 | 5 | 32 | 12.1 |
| A2 | B10 | C16 | D6 | 6 | 64 | 8 |
| A2 | B10 | C16 | D6 | 7 | 32 | 16 |
| A5 | B10 | C1 | D5 | 7 | 16 | 7.5 |
| A5 | B10 | C1 | D5 | 8 | 16 | 6.7 |
| A5 | B10 | C2 | D5 | 5 | 32 | 5.9 |
| A5 | B10 | C2 | D5 | 7 | 32 | 5.2 |
| A5 | B10 | C8 | D5 | 7 | 32 | 5.8 |
| A5 | B10 | C8 | D5 | 8 | 8 | 17.6 |
| A5 | B10 | C13 | D5 | 5 | 16 | 3.3 |
| A5 | B10 | C13 | D5 | 7 | 16 | 6.8 |
| A5 | B10 | C16 | D5 | 3 | 64 | 8 |
| A5 | B10 | C16 | D5 | 5 | 32 | 4.2 |
| A5 | B10 | C16 | D5 | 7 | 16 | 7.9 |
| A8 | B10 | C10 | D5 | 7 | 16 | 15.9 |
| A8 | B10 | C13 | D3 | 6 | 32 | 11.9 |
| A8 | B10 | C13 | D5 | 7 | 16 | 8.7 |
| A8 | B10 | C13 | D6 | 8 | 16 | 14.7 |
| A8 | B10 | C16 | D2 | 8 | 8 | 33.3 |
| A8 | B10 | C16 | D3 | 5 | 32 | 9.4 |
| A8 | B10 | C16 | D3 | 6 | 32 | 10.1 |
| A8 | B10 | C16 | D6 | 5 | 64 | 8 |

See Fig. S1 for the building block index and Fig. S2 for the ratio index.

**Table S5 | Summary of atom features.**

| Feature | Dimension | Description |
|---|---|---|
| Atomic number | 102 | The atomic number in the periodic table |
| Atom degree | 12 | The number of directly bonded atoms to a specific atom |
| Formal charge | 1 | The electric charge assigned to an atom |
| Num radical electrons | 6 | The number of unpaired electrons present on an atom |
| Hybridization | 6 | Mixing of atomic orbitals to form new hybrid orbitals |
| Aromatic | 1 | Whether the atom is part of an aromatic ring system |
| Total num H | 6 | The total number of hydrogen atoms attached to the atom |
| Chiral | 1 | Whether the atom is a chiral center |
| Chirality type | 2 | The chiral configurations: R/S |
| Atom mass | 1 | The atomic mass of an atom |

**Table S6 | Summary of bond features.**

| Feature | Dimension | Description |
|---|---|---|
| Bond type | 5 | The type of chemical bond, such as single, double, triple |
| Conjugation | 1 | Whether the bond is part of a conjugated system |
| In ring | 1 | Whether the bond is in a ring system |
| Stereo | 7 | The spatial arrangement of atoms around the bond |

**Table S7 | Hyperparameter setting and search range in base learner training stage.**

| Hyperparameters | Value |
|---|---|
| Learning rate | [5e-5, 1e-4, 2e-4, 3e-4, 5e-4] |
| Dropout | [0, 0.05, 0.1, 0.2] |
| Weight decay | [0, 1e-6, 1e-4] |
| Batch size | 32 |
| Patience | 20 |

**Table S8 | Hyperparameter setting in active learning stage.**

| Hyperparameters | Value | Description |
|---|---|---|
| n_estimators | 20 | The number of base learners |
| $\lambda_{\text{antibacterial}}$ | 0.7 | Weights for the antibacterial property prediction task |
| $\lambda_{\text{hemolytic}}$ | 0.3 | Weights for the hemolytic property prediction task |
| $\beta$ | 2 | Weights used to balance exploitation and exploration |
| $\gamma$ | 0.1 | Proportion of top-ranked samples for clustering |
| n_clusters | 20 | The number of clusters in K-means clustering |
| $k$ | 3 | The number of samples selected in each cluster |
